# Systematic functional analysis of Rab GTPases reveals limits of neuronal robustness in *Drosophila*

**DOI:** 10.1101/2020.02.21.959452

**Authors:** Friederike E. Kohrs, Ilsa-Maria Daumann, Bojana Pavlović, Eugene Jennifer Jin, Shih-Ching Lin, Fillip Port, F. Ridvan Kiral, Heike Wolfenberg, Thomas F. Mathejczyk, Chih-Chiang Chan, Michael Boutros, P. Robin Hiesinger

**Affiliations:** Division of Neurobiology, Institute for Biology, Freie Universität Berlin, Germany; German Cancer Research Center (DKFZ), Div. Signaling and Functional Genomics and Heidelberg University, Heidelberg, Germany; University of California, San Diego, La Jolla, CA, USA; National Taiwan University, Taipei, Taiwan

## Abstract

Rab GTPases are molecular switches that regulate membrane trafficking in all cells. Neurons have particular demands on membrane trafficking and express numerous Rab GTPases of unknown function. Here we report the generation and characterization of molecularly defined null mutants for all 26 *rab* genes in *Drosophila*. In addition, we created a transgenic fly collection for the acute, synchronous release system RUSH for all 26 Rabs. In flies, all *rab* genes are expressed in the nervous system where at least half exhibit particularly high levels compared to other tissues. Surprisingly, loss of any of these 13 nervous-system enriched Rabs yields viable and fertile flies without obvious morphological defects. However, 9 of these 13 affect either developmental timing when challenged with different temperatures, or neuronal function when challenged with continuous stimulation. These defects are non-lethal under laboratory conditions, but represent sensitized genetic backgrounds that reveal limits of developmental and functional robustness to environmental challenges. Interestingly, the neuronal *rab26* was previously proposed to function in synaptic maintenance by linking autophagy and synaptic vesicle recycling and we identified *rab26* as one of six *rab* mutants with reduced synaptic function under continuous stimulation conditions. However, we found no changes to autophagy or synaptic vesicle markers in the *rab26* mutant, but instead a cell-specific role in membrane receptor turnover associated with cholinergic synapses in the fly visual system. Our systematic functional analyses suggest that several Rabs ensure robust development and function under varying environmental conditions. The mutant and transgenic fly collections generated in this study provide a basis for further studies of Rabs during development and homeostasis *in vivo*.

## INTRODUCTION

Rab GTPases have been named for their initial discovery in brain tissue (<u>Ra</u>s-like proteins from rat <u>b</u>rain), where their abundance and diversity reflect neuronal adaptations and specialized membrane trafficking (Kiral et al., 2018; Touchot et al., 1987). Yet, Rabs are found in all eukaryotic cells, where they function as key regulators of membrane trafficking between various membrane compartments (Pfeffer, 2017; Zhen and Stenmark, 2015). Consequently, Rab GTPases are commonly used as markers, and some have become gold standard identifiers of various organelles and vesicles in endocytic and secretory systems(Pfeffer, 2017; Zerial and McBride, 2001).

Over the years, Rab GTPases have repeatedly been analyzed as a gene family to gain insight into membrane trafficking networks (Best and Leptin, 2020; Chan et al., 2011; Dunst et al., 2015; Gillingham et al., 2014; Gurkan et al., 2005; Jin et al., 2012; Pfeffer, 1994; Stenmark, 2009; Zerial and McBride, 2001). However, a complete and comparative null mutant analysis of all family members is not available for any multicellular organism. The *Drosophila* genome contains 31 potential *rab* or rab-related genes, of which 26 have been confirmed to encode protein-coding genes (Chan et al., 2011; Dunst et al., 2015; Jin et al., 2012), compared to 66 *rab* genes in humans (Gillingham et al., 2014). Of the 26 *Drosophila rab* genes, 23 have direct orthologs in humans that are at least 50% identical at the protein level, indicating high evolutionary conservation (Chan et al., 2011; Zhang et al., 2007).

In the nervous system, Rab GTPases have been predominantly associated with functional maintenance and neurodegeneration (Kiral et al., 2018; Veleri et al., 2018). For example, mutations in *rab7* cause the neuropathy CMT2B (Cherry et al., 2013; Spinosa et al., 2008; Verhoeven et al., 2003), Rab10 and other Rabs are phosphorylation targets of the Parkinson’s Disease-associated kinase LRRK2 (Dhekne et al., 2018; Steger et al., 2017), and Rab26 and Rab35 have been implicated in synaptic vesicle recycling (Binotti et al., 2015; Sheehan et al., 2016; Uytterhoeven et al., 2011). Neuronal longevity and morphological complexity have been suggested to require specific Rab-mediated membrane trafficking (Jin et al., 2018a; Jin et al., 2018b), but the roles of most Rab GTPases in neuronal development and function remain unknown.

We have previously developed a transgenic *Drosophila rab*-Gal4 collection based on large genomic fragments and a design for subsequent homologous recombination to generate molecularly defined null mutants (Chan et al., 2011; Jin et al., 2012). Analyses of the cellular expression pattern and subcellular localization based on YFP-Rab expression under its endogenous regulatory elements by us and others (Dunst et al., 2015) have revealed numerous neuronal Rabs with synaptic localization (Chan et al., 2011). We originally found that all 26 *Drosophila* Rab GTPases are expressed somewhere in the nervous system and half of all Rabs are enriched or strongly enriched in neurons (Chan et al., 2011; Jin et al., 2012). A more recent collection of endogenous knock-ins identified more varied expression patterns when more tissues were analyzed, but also validated the widespread neuronal and synaptic expression (Dunst et al., 2015). The function of most Rabs with strong expression in the nervous system has remained unknown.

Here we provide the first complete null mutant comparative analysis of all *rab* genes in a multicellular organism. We find that viability, development and neuronal function are highly dependent on environmental conditions in these mutants. Under laboratory conditions with minimal selection pressure, seven mutants are lethal, one semi-lethal with few escapers, two are infertile and six are unhealthy based on progeny counts. All 13 nervous-system enriched Rabs are viable under laboratory conditions, but 9 of these 13 exhibit distinct developmental or functional defects depending on environmental challenges. Our survey and fly collection provide a basis to systematically elucidate Rab-dependent membrane trafficking underlying development and function of all tissues in a multicellular organism.

## RESULTS

### Generation of the *rab* GTPase null mutant collection

Our earlier observation of a synaptic localization of all nervous-system-enriched Rabs led us to speculate that many Rab GTPases may serve roles related to neuron-specific functions. To test this idea, we set out to generate a complete null mutant collection. We have previously published molecularly defined null mutants of *rab27* (Chan et al., 2011) and *rab7* (Cherry et al., 2013) as Gal4 knock-ins using a BAC recombineering/homologous recombination approach (Chan et al., 2011). Seven additional molecularly defined null mutants have previously been reported: *rab1* (Thibault et al., 2004), *rab3* (Graf et al., 2009), *rab5* (Wucherpfennig et al., 2003), *rab6* (Purcell and Artavanis-Tsakonas, 1999), *rab8* (Giagtzoglou et al., 2012), *rab11* (Bellen et al., 2004) and *rab32* (Ma et al., 2004). For the remaining 17 *rab* genes we generated six null mutants as Gal4 knock-ins that replace the endogenous open reading frames, or the ATG start codon, using homologous recombination; these include *rab2, rab4, rab19, rab30, rabX1* and *rabX6* (Fig. 1A; Suppl. Fig. 1A-B). The remaining 11 null mutants were generated using CRISPR/Cas9, including *rab9, rab10, rab14, rab18, rab21, rab23, rab26, rab35, rab39, rab40*, and *rabX4* (Fig. 1A; Suppl. Fig. 1C-D). All mutants were molecularly validated as described in detail in the Material and Methods.

**Figure 1:**
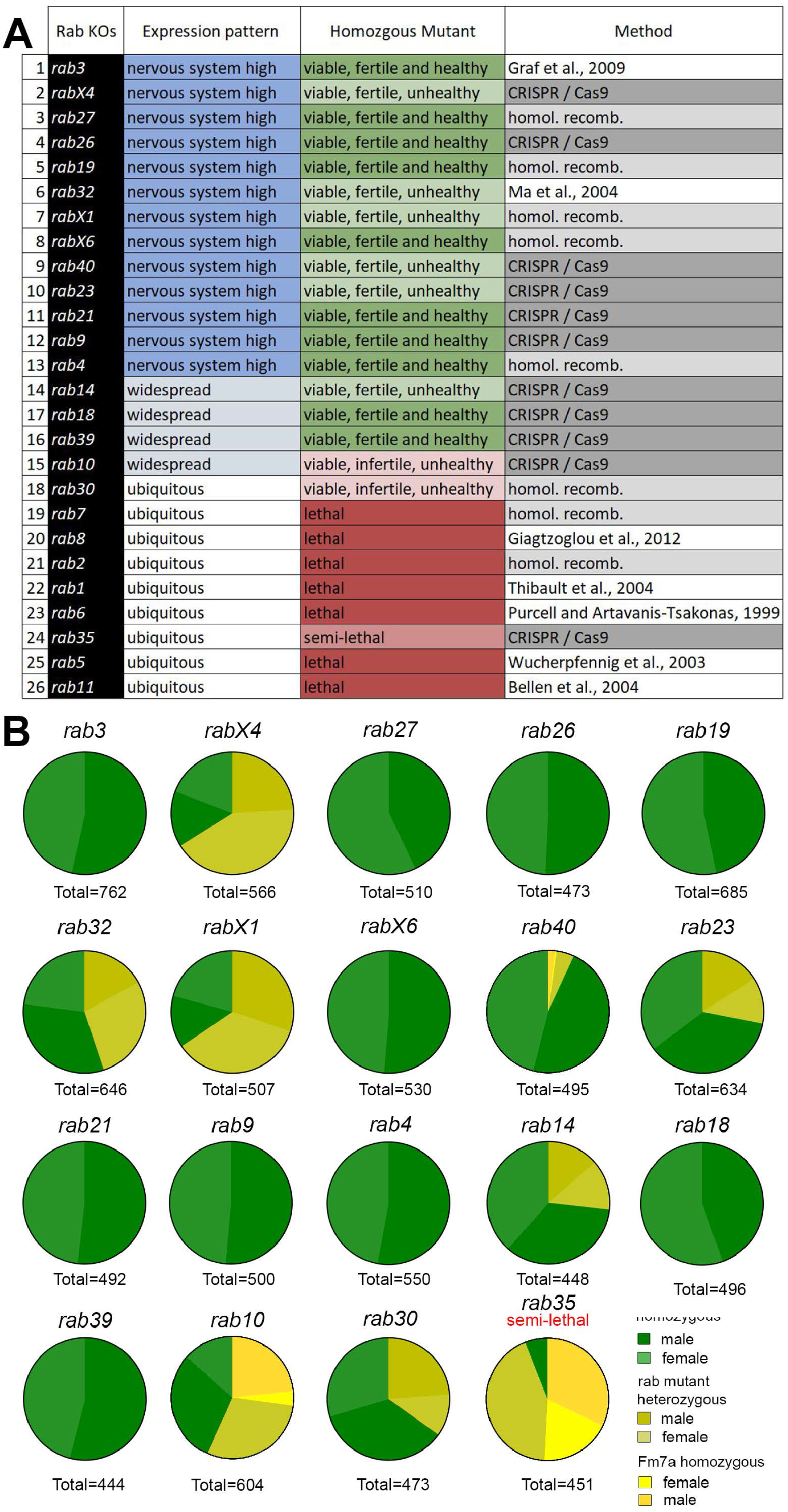
Generation and viability analysis of a complete *rab* null mutant collection. (A) List of all 26 existing *Drosophila rab* null mutants, sorted by their expression pattern from nervous system high to ubiquitous. Two thirds of the *rab* mutants are homozygous viable and fertile. 8 *rab* mutants are lethal when homozygous. The method used for mutant generation is indicated. (B) Pie charts show female/male (shades of green) distribution of homozygous viable *rab* mutants and their ratio of homozygosity after ten generations. Ten of the 18 viable or semi-lethal *rab* mutants are fully homozygous, while the others still retain their balancer chromosome (shades of yellow) to varying degrees. Total numbers of flies counted are indicated in the figure.

### All nervous system-enriched *rab* mutants are viable under laboratory conditions

All mutant chromosomes were tested for adult lethality in homozygosity. Of the 26 null mutants, seven are homozygous lethal *(rabs 1, 2, 5, 6, 7, 8, 11)* and one, *rab35*, is homozygous semilethal with few male escapers; 18 *rab* null mutants are viable as homozygous adults under laboratory conditions (Fig. 1A).

All mutants were initially generated with the null mutant chromosome in heterozygosity over a balancer chromosome. Balancers contain multiple genetic aberrations, rendering them generally less healthy than wild type chromosomes; balancer chromosomes are therefore outcompeted in healthy stocks after a few generations. However, after 10 generations, only 10 of the 18 viable lines lost the balancer, indicating that eight *rab* mutant chromosomes confer a competitive disadvantage (Fig. 1B). For five *rab* mutant chromosomes *(rab14, rab23, rab30, rab32*, and *rab40)* a minority of balanced flies remained in the viable stocks, suggesting that the mutant chromosomes conferred mildly reduced viability in homozygosity. For *rab10, rabX1*, and *rabX4* balanced mutant flies were in the majority, indicating substantially disadvantageous mutant chromosomes (Fig. 1B). Sibling crosses between unbalanced homozygous mutant flies revealed an inability to lay eggs for *rab10* mutant flies and only non-developing eggs for *rab30* mutant flies, which was rescued by overexpression with the *rab30*-Gal4 line. In all other cases, homozygous mutant eggs developed, albeit in some cases at significantly lower numbers or at altered developmental speeds, as discussed in detail below. These observations suggest a range of mutant effects that may affect development or function, yet remain sub-threshold for viability under laboratory conditions.

Surprisingly, all lethal mutants exhibit ubiquitous expression, while all 13 Rab GTPases that we previously reported to be enriched in the nervous system are viable and fertile (Fig. 1A). This observation once again put a spotlight on the question of specialized Rab GTPase functions in the nervous system. The development and maintenance of the nervous system requires robustness to variable and challenging conditions, including temperature and intense stimulation. We therefore hypothesized that many Rabs may provide context-specific neuronal roles that ensure robust development and function that are not apparent under laboratory rearing conditions. We therefore devised a series of assays to test all 16 viable and fertile *rab* null mutant stocks for development, function and maintenance under different challenging conditions.

### The majority of viable *rab* mutants affect developmental robustness at different temperatures

First, we analyzed developmental robustness to temperatures of 18°C, 25°C and 29°C (Fig. 2A-C). The 16 homozygous viable and fertile mutants include all nervous system-enriched *rabs* plus *rab14, rab18* and *rab39*. At 25°C twelve of these 16 homozygous stocks produced wild type numbers of offspring within 10-11 days comparable to control (Fig. 2B; Suppl. Table 1). *rab19, rab40, rabX1* and *rabX4* all exhibited significantly delayed development. *rabX1* mutant flies laid very few eggs, and only few of those eclosed as adults (20% of control). By contrast, *rabX4* mutant flies laid a high number of eggs, but most of these did not develop; only few *rabX4* adult escapers developed with 2-4 days developmental delay (Fig. 2B; Suppl. Table 1).

**Figure 2:**
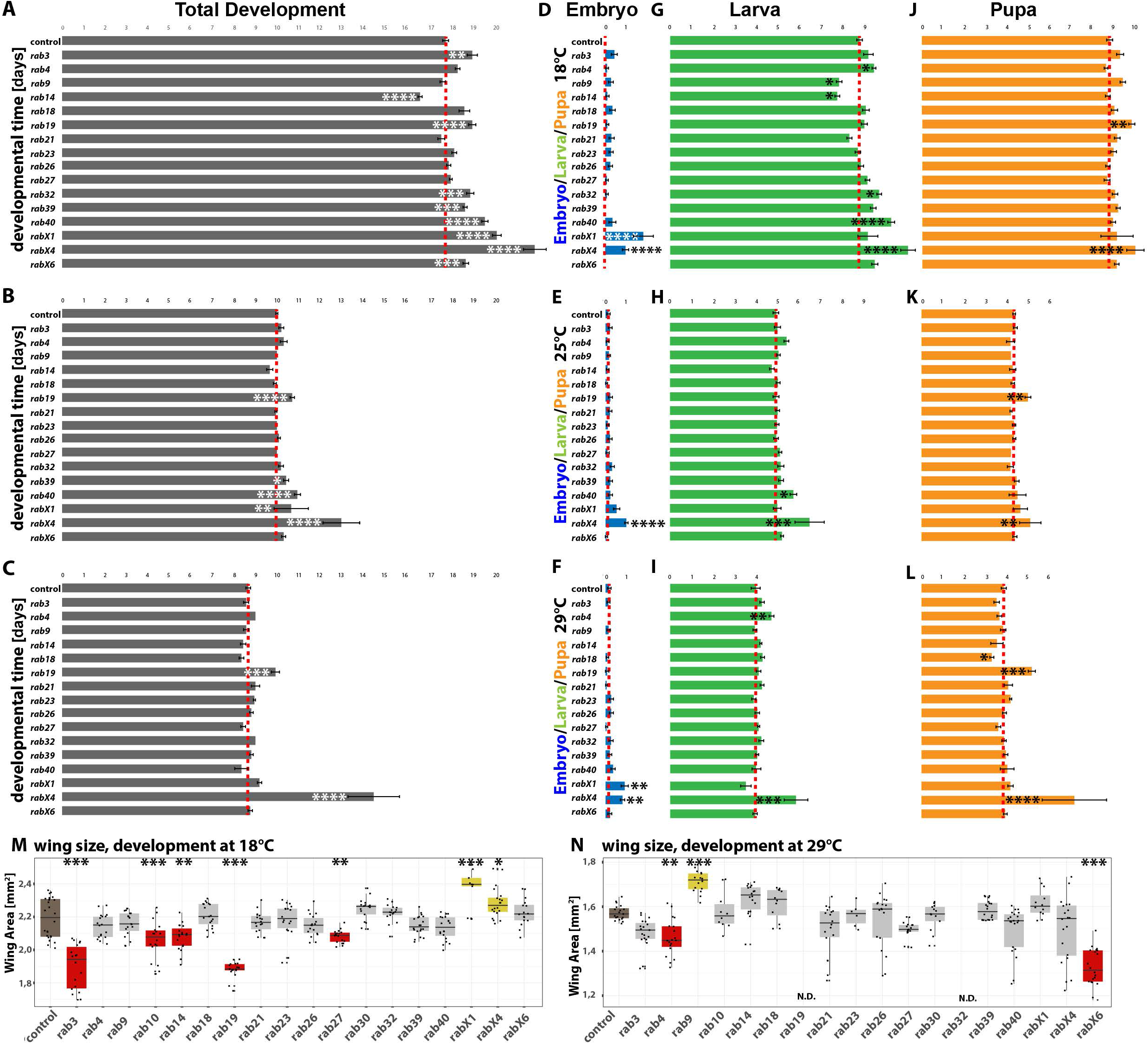
Developmental analyses of all viable *rab* mutants at different temperatures. (A-C) Total developmental time at 18°C (A), 25°C (B), and 29°C (C) for all homozygous viable *rab* mutants. (D, G, and J) Developmental time at 18°C for all homozygous viable *rab* mutants, separated into embryonal (blue, D), larval (green, G) and pupal (orange, J) phases. (E, H, and K) Developmental time at 25°C for all homozygous viable *rab* mutants, separated into embryonal (blue, E), larval (green, H) and pupal (orange, K) phases. (F, I, and L) Developmental time at 29°C for all homozygous viable *rab* mutants, separated into embryonal (blue, F), larval (green, I) and pupal (orange, L) phases. Mean ± SEM; *p < 0.05, **p < 0.01, ***p < 0.001, ****p < 0.0001; Unpaired non-parametric Kolmogorov-Smirnov test. (M- N) Wing surface area measurement for all homozygous viable *rab* mutants at 18°C (M) and 29°C (N). Wild type (brown) and *rab* mutant (grey) wing size. Significantly reduced (red) and increased wing sizes (yellow) compared to control are highlighted. Boxplot with horizontal line representing the median; individual data points are represented as dots. 18-22 wings per genotype were quantified; *p < 0.05, **p < 0.01, ***p < 0.001, ****p < 0.0001; Ordinary one-way ANOVA with pair-wise comparison, Tukey HSD test.

Development at 29°C revealed aberrations in three of these four mutants (Fig. 2C; Suppl. Table 1). *rabX4* had very few escapers (more than 100-fold reduced viability) with the longest developmental delay amongst mutants. *rabX1* exhibited a smaller delay with significantly reduced numbers of eclosing adults (Suppl. Table 1). *rab19* exhibited a significant developmental delay and a 50-80% rate of late pupal lethality at 29°C, that was not observed at lower temperatures. All *rab19* adults raised at 29°C died within a few days. Finally, *rab32* also exhibited increased late pupal lethality specifically at 29°C, while survivors exhibited no developmental delay and normal eclosion times. These findings suggest a particular sensitivity to higher temperatures for *rab19* and *rab32*, as well as reduced viability at all temperatures for *rabX1* and *rabX4*. Development at 18°C revealed increased variability and developmental delays for 8 of the 16 mutants (Fig. 2A; Suppl. Table 1). Reduced viability at 18°C was observed only for *rabX1* and *rabX4*.

Our observations raised the question whether developmental delays were present during specific developmental stages. We therefore performed the developmental timing assay separately for embryo, larval and pupal development (Fig. 2D-L). We collected embryos after a 24 hours egg-laying period and measured hatching times of the first 1st instar larvae (Fig. 2D-F), the first larvae transitioning to pupae (Fig. 2G-I), and the first adults to eclose (Fig. 2J-L) at all three temperatures.

Hatching times of 1st instar larvae were similar in all mutants except two (*rabX1* and *rabX4;* Fig. 2D-F), consistent with previous reports of robust temporal compensation of embryogenesis (Chong et al., 2018). Larval development at 18°C lasted on average a little less than twice as long compared to 25°C and timing was more variable between mutants (Fig. 2G, H); pupal development lasted almost exactly twice as long in all mutants (Fig. 2J, K). At 29°C both larval and pupal developmental timing was further sped up by 15-20% on average compared to 25°C (Fig. 2I, L).

Significant delays of embryo development were observed for *rabX1* and *rabX4* at all temperatures (Fig. 2D-F). By contrast, larval development at 25°C or 29°C was only affected for *rab4, rab40* and *rabX4*. At 18°C variability of larval development amongst mutants was higher and *rab32* also exhibited delayed development, while *rab9* and *rab14* exhibited shortened larval periods (Fig. 2G-I).

Pupal development did not exhibit an increased variability amongst mutants, in contrast to the variability observed for larval stages. Pupal development was only increased in two mutants, including *rabX4*, which exhibited delays and reduced viability at all temperatures and developmental stages. The second, *rab19*, was the only mutant that caused a developmental delay specific to pupal stages, as embryo and larval development were similar to control; *rab19* viability was not affected at at 18°C and 25°C, in contrast to *rab19’s* highly increased sensitivity to 29°C. Conversely, pupal development was not affected at any temperature for the *rab9, rab14, rab32* and *rab40* mutants that exhibited altered larval periods.

Temperature is known to affect organ development through changes in cell size (Azevedo et al., 2002). For example, the *Drosophila* wing in control flies is 20-30% larger after development at 18°C compared to development at 29°C, as previously shown (Fig. 2M, N). As for overall developmental time at 18°C, we found wing sizes to be highly variable between mutants at 18°C (Fig. 2M; Suppl. Fig. 2A-B). We observed significantly smaller wings for *rab3, rab10, rab14, rab19* and *rab27*. By contrast, the two mutants with the longest developmental delay at 18°C, *rabX1* and *rabX4*, exhibited significantly increased wing size (Fig. 2M-N; Suppl. Fig. 2C, E). At 29°C, wings were significantly smaller in all mutants (Fig. 2N). The smallest reduction (largest wing size) at 29°C was observed in the *rab9* mutant and significant reductions were observed in *rab4* and *rabX6* mutants (Fig. 2M-N; Suppl. Fig. 2F). Our *rab23* null mutant exhibited the planar cell polarity phenotype of wing bristles reported previously (Dunst et al., 2015; Pataki et al., 2010). The temperature-dependent changes in wing size were similar to control in *rab23* mutant wings. In addition, we observed a previously not reported highly penetrant transversal p-cv vein shortening (in 90% of the wings studied) at 18°C, which was ameliorated at 29°C (12% penetrance) (Suppl. Fig. 2G-J). In sum, all mutants that affect viability or development at different temperatures also exhibited altered wing development.

We conclude that 8 of the 13 null mutants of Rabs previously found to be strongly expressed in the nervous system exhibit various defects in developmental timing (Table 1): *rabX1* affected viability and specifically embryo development, but timing remains robust at different temperatures; *rab4* and *rab40* specifically affect larval development; *rab19* specifically affects pupal development, an effect that is again most pronounced at high temperature. Finally, *rabX4* affected all developmental stages in a heat-sensitive manner. Developmental robustness to temperature was affected in mutants for *rabX4* (all stages), *rab19* (delayed pupae, pupal lethality), *rab32* (pupal lethality), *rab9* and *rab14* (reduced larval period). In addition to these eight, mutants for *rab3, rab39* or *rabX6* exhibited overall developmental delays at 18°C (Fig. 2A), but no significant differences when analyzed separately for embryo, larval and pupal development.

**Table 1:**
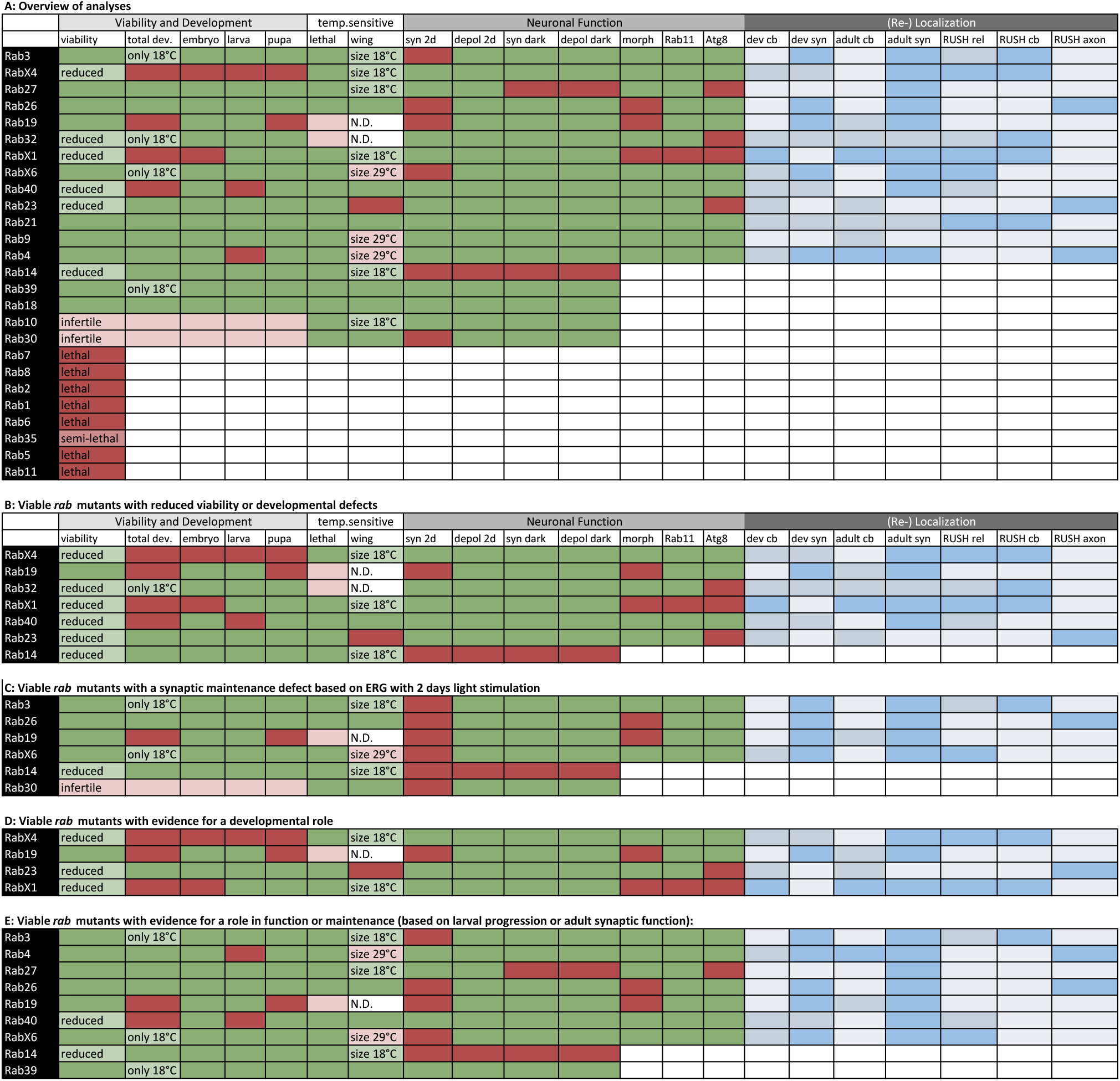
Summary of functional and RUSH analyses. (A) Overview of analyses (‘Viability and Development’, ‘temp. sensitive’, ‘Neuronal Function’, and ‘(Re-) Localization’) done in this study for all Rab GTPases. (B) List of all viable *rab* mutants, which show a reduced viability or developmental defects. (C) List of all viable *rab* mutants, which show synaptic maintenance defects after 2 days light stimulation based on ERG recordings. (D) List of all viable *rab* mutants, which show evidence for a developmental role. (E) List of all viable *rab* mutants, which show evidence for a role in function or maintenance. Abbreviations: cb = cell body, depol = depolarization, dev. = development, morph = morphology, N.D. = not determined, Rel = release, syn = synaptic, 2d = 2 days. Color code: lighter green to darker red denotes increasing deviation from controls in functional analyses. Darker to lighter blue indicates higher to lower localization or relocalization of Rab proteins.

### Several viable *rab* mutants affect the maintenance of stimulus-dependent synaptic function

To specifically challenge neuronal function and maintenance, we tested the effect of light stimulation of photoreceptor neurons, a widely used model to identify mutants affecting neuronal maintenance and degeneration in *Drosophila* (Jaiswal et al., 2012). To quantitatively analyze alterations of photoreceptor response properties to light stimulations, we established a protocol based on sensitization curves for electroretinogram (ERG) recordings of photoreceptor neurons. The ERG is an extracellular recording that reveals two aspects of neuronal function: first, the depolarization measures the ability of photoreceptor neurons to convert a light stimulus into an electrical signal. Reduced depolarization can be the results of a reduced ability to perceive light (phototransduction), reduced electrical properties of individual cells, or loss of neurons. Second, the ERG indicates the ability to transmit the presynaptic signal to the postsynaptic interneurons via the so-called ‘on’ transient. Loss of the ‘on’ transient can result from defective neurotransmission or degeneration that starts at the synapse, as shown for the *rab7* mutant previously (Cherry et al., 2013). The ERG is mostly used as a qualitative method, because both depolarization and ‘on’ transient intensities are highly sensitive to differences in genetic background, eye pigmentation, intensity of the light stimulus and other recording variables. To identify a sensitive period where mild alterations of neuronal function and maintenance should be discernible within the dynamic range of the measurement, we established sensitization curves over several days of stimulation. In control flies, continuous stimulation lead to a gradual decline of ‘on’ transient amplitude (Fig. 3A) and depolarization (Fig. 3B) over a 7-days period. Two days light stimulation represent a highly sensitized period with a dynamic range for improvement or worsening of potential defects for both the ‘on’ transient (Fig. 3A) and depolarization (Fig. 3B).

**Figure 3:**
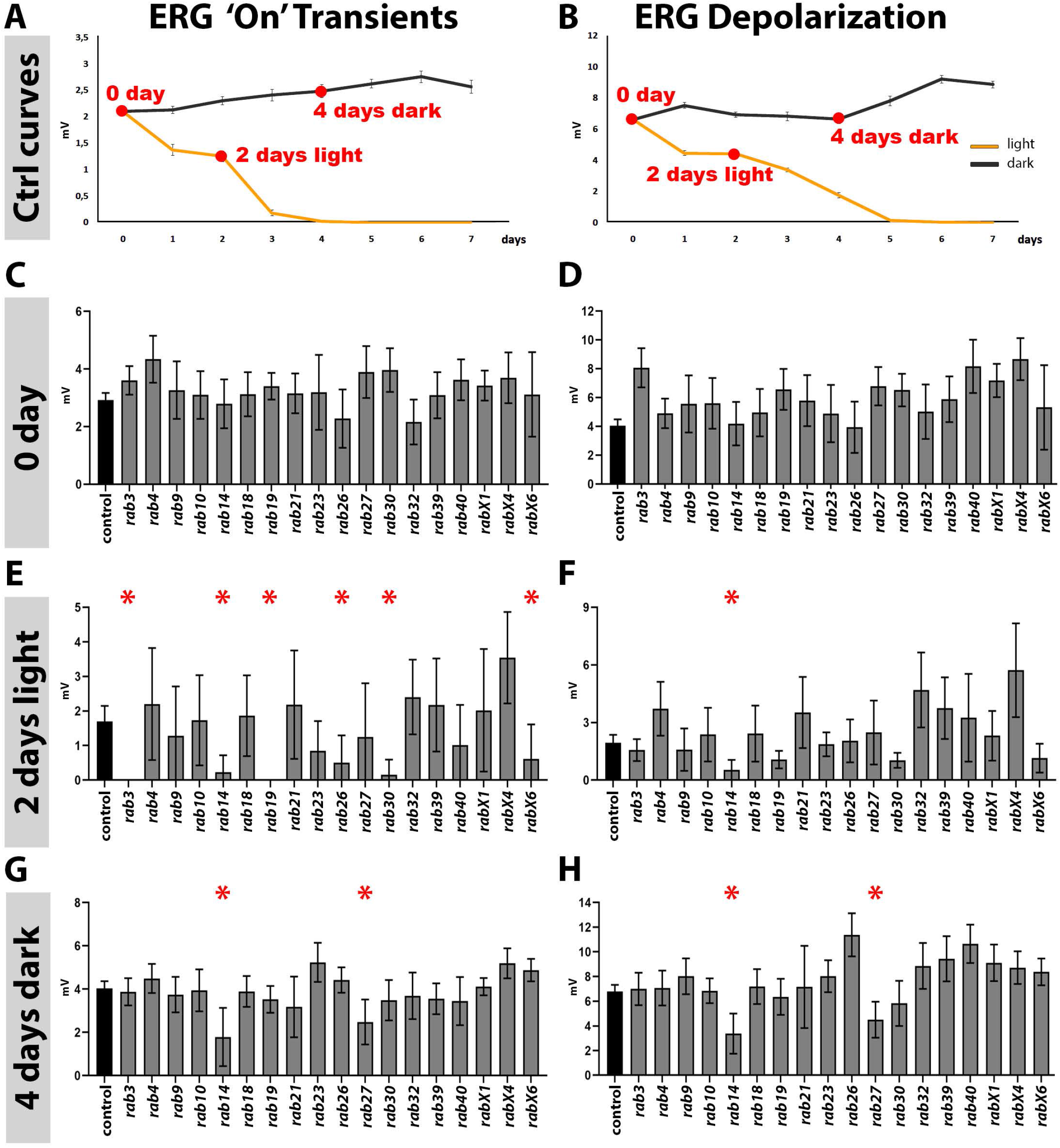
Analysis of neuronal function and maintenance based on electroretinograms. (A-B) Sensitization curves for light stimulated (orange curve) and dark-reared (black curve) wild type flies generated by electroretinogram (ERG) recordings. ‘on’ transient signal is lost after 4 days of light stimulation. Complete loss of depolarization signal after 5 days of light stimulation. 0 day, 2 days light stimulation and 4 days dark-rearing are highlighted in red. Mean ± SEM; 25-30 flies were recorded for each day (0-7 days) and each condition (light and dark); Ordinary one-way ANOVA with pair-wise comparison. (C-D) ‘on’ transient and depolarization of newly hatched (0 day) flies. Wild type control in black, all homozygous viable rab mutants in grey. (E-F) ‘on’ transient and depolarization of wild type (black) and homozygous viable *rab* mutants (grey) after 2 days of light stimulation. (G-H) ‘on’ transient and depolarization of wild type (black) and homozygous viable *rab* mutants (grey) after 4 days of dark-rearing. (C-H) Mean ± SD; *p < 0.05; 25-30 flies were recorded for each genotype and condition; Ordinary one-way ANOVA with group-wise comparison.

For all 18 viable *rabs*, we tested mutants in a *white minus* background. All mutants further expressed the *3xP3-RFP* marker with the exception of existing mutations in *rab3* and *rab32*. These two mutants were previously generated in different genetic backgrounds (Graf et al., 2009; Ma et al., 2004) and we crossed these mutant chromosomes in *white minus* backgrounds for all experiments in our study.

First, we performed ERG recordings of newly hatched flies to assess neuronal function immediately after development (‘0 day’; Fig. 3A-D). None of the mutants exhibited significant reductions of their ‘on’ transient (Fig. 3C) or depolarization (Fig. 3D) immediately after hatching (0 day). Next, we used continuous light stimulation to measure alterations to function in a stimulus-dependent manner (Fig. 3E-H) and dark-rearing to assess alterations to photoreceptor function in an age-dependent, stimulus-independent manner (Fig. 3G-H).

After two days of light stimulation, six *rab* mutants exhibited significantly reduced neurotransmission compared to control based on their ‘on’ transients: *rab3, rab14, rab19, rab26, rab30* and *rabX6*. Of these six mutants, only *rab14* exhibited a significantly decreased depolarization, indicating reduced cellular function. *rab14* was also the only Rab that was not in our original list of ‘neuron-enriched Rabs’, while the other five encode synaptic proteins. By contrast to these loss-of-function phenotypes, *rabX4* exhibited significantly increased ‘on’ transients and depolarization, despite its developmental delay and reduced viability. This observation is not easily explained, but suggests increased sensitization of synaptic transmission to the stimulus.

To test whether these maintenance defects were strictly stimulus-dependent, we compared these results to 2-days and 4-days dark-rearing. *rab14* exhibited similar decreases in both ‘on’ transient and depolarization after four days in the dark, suggesting stimulus-independent functional defects. By contrast, none of the other five neuron-enriched Rabs *(rab3, rab19, rab26, rab30* and *rabX6)* exhibited reduced neurotransmission or health based on ERGs in the absence of a light stimulus. Finally, *rab27* exhibited a defect in both ‘on’ transient and depolarization specifically after four days in the dark, but not after light-stimulation. Together, these findings suggest stimulus-dependent synaptic defects for five neuron-enriched Rabs *(rab3, rab19, rab26, rab30* and *rabX6)*, alterations of both synaptic and other cellular functions for three Rabs, the neuron-enriched *rab27* and *rabX4*, as well as for the ubiquitous *rab14*.

Next, we analyzed the morphology of photoreceptor axon projections after light stimulation compared to newly hatched flies using an antibody against the photoreceptor membrane protein Chaoptin. All 13 nervous system-enriched *rab* mutants exhibited axonal projections that were indistinguishable from control in newly hatched flies (Suppl. Fig. 3). Amongst newly hatched flies, only *rabX1* exhibited strongly increased accumulations of Chaoptin in non-photoreceptor cell bodies surrounding the neuropils (Fig. 4A), a phenotype previously observed for endomembrane degradation mutants including *rab7* (Cherry et al., 2013) and the *v-ATPase v100* (Williamson et al., 2010).

**Figure 4:**
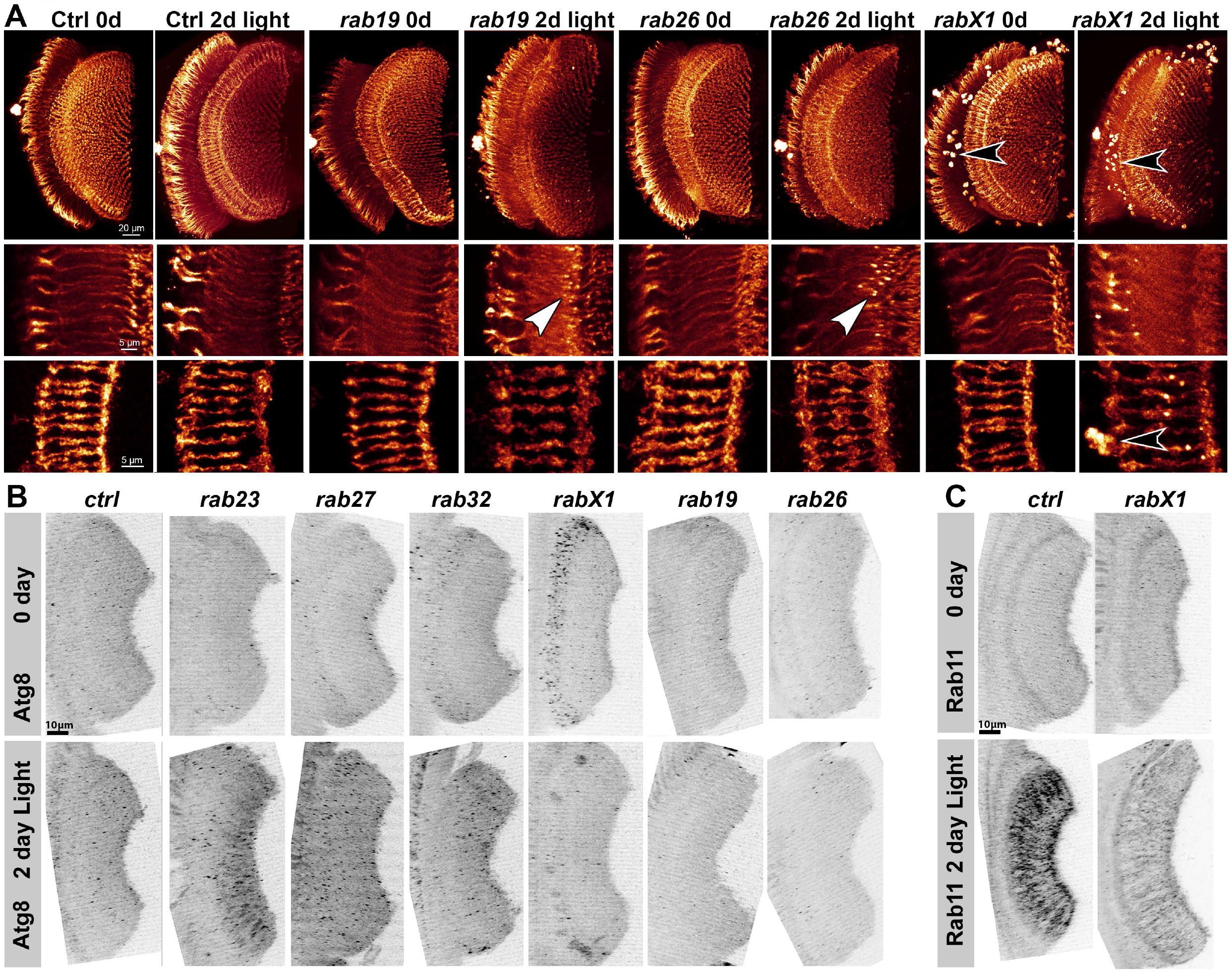
Analyses of morphology, recycling endosomal function (Rab11) and autophagy (Atg8) at photoreceptor axon terminals after continuous light stimulation. (A) Chaoptin-labeling of 0 day and 2 days light stimulated wild type and *rab* mutant photoreceptor projections (overview top panel, R1-R6 middle panel, R7-R8 bottom panel). The *rabX1* mutant exhibits Chaoptin accumulations in non-photoreceptor cell bodies independent of stimulation (black arrowheads). After 2 days of light stimulation, *rab26* and *rab19* mutants display membrane accumulations in their axon terminals (white arrowheads). Scale bar: 20 μm (top panel), 5 μm (middle and bottom panel); number of brains n = 3-5 per antibody staining. (B) Labeling of photoreceptor projections in retina-lamina preparations against the autophagosome marker Atg8 of newly hatched and 2 days light stimulated wild type and six *rab* mutants. Only *rab23, rab27*, and *rab32* show significant increases in Atg8-positive compartments after 2 days light stimulation. Scale bar: 10 μm; number of retina-lamina preparations n = 3 for each condition and staining. (C) Labeling of photoreceptor projections in retina-lamina preparations against the recycling endosome marker Rab11 of newly hatched and 2 days light stimulated wild type and *rabX1* flies. Suppressed increase in Rab11 levels after 2 days of light stimulation in *rabX1*. Scale bar = 10 μm; number of retina-lamina preparations n = 3 for each condition and staining.

After light stimulation, two mutant exhibited alterations of their axon terminal morphology. Mutants for *rab26*, and to a lesser extent *rab19*, exhibited distinct membrane accumulations at the distal tips of R1-R6 photoreceptor axon terminals (Fig. 4A).

We next asked whether autophagosome formation was affected in *rab* mutant axon terminals. Atg8/LC3-positive autophagosomes were relatively infrequent given the number of axon terminals in the lamina and only mildly more frequent after light stimulation (Fig. 4B). Significant increases of Atg8-positive compartments in the lamina were observed for *rab23, rab27* and *rab32*, as well as in some cell bodies for *rabX1* (Fig. 4B). Notably, none of these mutants exhibited functional defects after light stimulation (comp. Fig. 3E-F; Table 1).

We previously showed that most nervous-system enriched Rabs encode proteins that colocalize with the recycling endosome marker Rab11 at photoreceptor axon terminals (Chan et al., 2011). After light stimulation, Rab11 is strongly upregulated in the synaptic terminals of wild type photoreceptor neurons as well as most mutants. By contrast, in *rabX1* mutant axon terminals, the increase in Rab11 levels was suppressed, suggesting a local defect in the formation of recycling endosomes, consistent with a recent report of a link between early and recycling endosomal function (Woichansky et al., 2016) (Fig. 4C). Surprisingly, the six *rab* mutants that specifically affected synaptic functional maintenance after light stimulation did not exhibit significant changes to their Rab11 or Atg8 levels at synaptic terminals. Hence, the viable Rabs implicated in synaptic functional maintenance might employ mechanisms different from Rab11-dependent endomembrane recycling and Atg8-dependent autophagy in fly photoreceptor neurons.

### Dynamic localization of nervous system-enriched Rabs: Establishing a toolbox to use the RUSH system for Rabs

To facilitate the analysis of intracellular localization dynamics of Rab GTPases we established the RUSH system for Rabs in *Drosophila*. RUSH stands for ‘<u>r</u>etention <u>u</u>sing <u>s</u>electable <u>h</u>ooks’ and is designed to allow the acute and synchronous release of tagged proteins from a ‘hook compartment’ (Fig. 5A) (Boncompain et al., 2012). The rationale for these experiments was to follow acutely released proteins as they synchronously change their localization. We have previously characterized the compartment-identities of constitutively expressed YFP-tagged Rabs in photoreceptor cell bodies and at synapses (Chan et al., 2011; Jin et al., 2012). Acute release has the potential to reveal which Rabs target specific compartments or localize diffusely in the cytoplasm.

**Figure 5:**
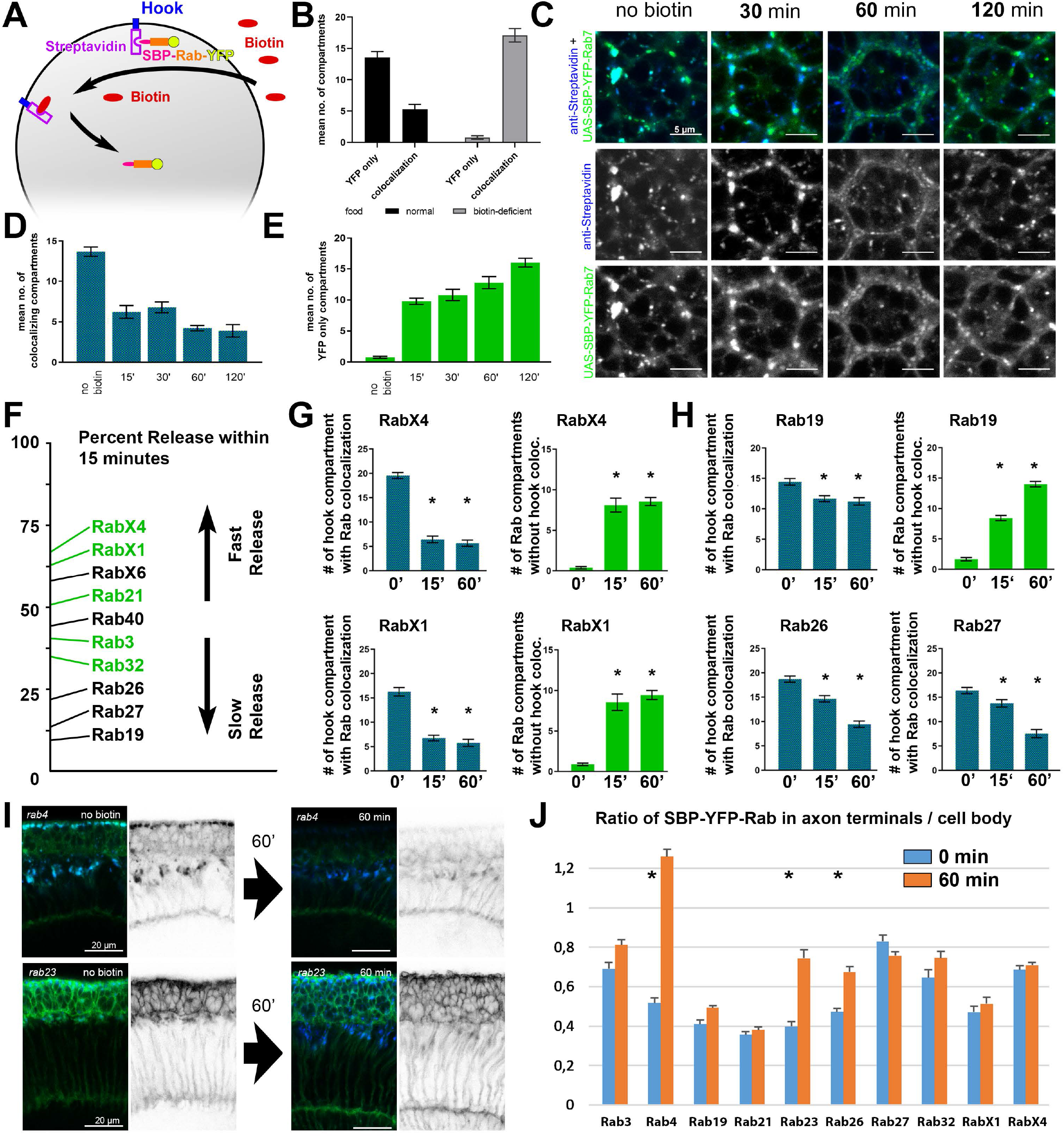
Establishment of the RUSH system for *Drosophila* Rab GTPases. (A) Schematic of the RUSH system. (B) Amount of retained RUSH-Rab7 (co-localization with hook) is significantly higher when raised on biotin-deficient compared to normal food. Amount of released RUSH-Rab7 (YFP only) is significantly higher when raised on normal food compared to biotin-deficient food. Mean ± SEM; **** p < 0.0001; number of ommatidia n = 9 for each condition (normal vs. biotin-def. food) from 3-4 animals; Ordinary one-way ANOVA with pair-wise comparison. (C) Biotin-release timeline shows separation of RUSH-Rab7 from the Golgi-hook and formation of Rab7-positive rings within 30-60 minutes. Scale bar = 5 μm. (D) Number of RUSH-Rab7 compartments co-localizing with the Golgi-hook is decreasing with biotin-incubation. Mean ± SEM; all **** p < 0.0001 compared to control; number of ommatidia n = 9 from 3-5 animals for each incubation time point; Ordinary one-way ANOVA with pair-wise comparison. (E) Number of RUSH-Rab7 compartments released from the Golgi-hook is increasing with biotin-incubation with biotin. Mean ± SEM; all **** p < 0.0001 compared to control; number of ommatidia n = 9 from 3-5 animals for each incubation time point; Ordinary one-way ANOVA with pair-wise comparison. (F) Percent release within 15 minutes of biotin-incubation for ten ‘nervous system high’ RUSH-Rabs. Most ‘fast-releasing’ Rabs display distinct compartments after release from the Golgi-hook (marked in green), while most ‘slow-releasing’ Rabs are diffusely localized in the cytoplasm. (G) RUSH-RabX1 and RUSH-RabX4 show ‘fast release’ behavior, more than 50% of the Rab-compartments are released from the Golgi-hook within the first 15 minutes of biotin-incubation. Mean ± SEM; *p < 0.05; number of ommatidia n = 9 from 3-5 animals for each incubation time point; Ordinary one-way ANOVA with pair-wise comparison. (H) RUSH-Rab19, RUSH-Rab26 and RUSH-Rab27 show ‘slow release’ behavior, less than 25% of the Rab-compartments are released from the Golgi-hook within the first 15 minutes of biotin-incubation. Mean ± SEM; *p < 0.05; number of ommatidia n = 9 from 3-5 animals for each incubation time point; Ordinary one-way ANOVA with pair-wise comparison. (I) Retina to axon terminal cross-sections reveal a significant increase in Rab-localization to the terminals after 60 minutes of biotin-incubation for RUSH-Rab4 and RUSH-Rab23. Single channel shows SBP-YFP-Rab localization. Scale bar = 20 μm. (J) Measurement of axon terminal to cell body fluorescence ratios reveals an increase in of axon terminal localization for RUSH-Rab4, RUSH-Rab23 and RUSH-Rab26. Mean ± SEM; *p < 0.05; Ordinary one-way ANOVA with pair-wise comparison.

First, we designed a series of four transgenic fly strains expressing a Streptavidin-hook, three for the endoplasmic reticulum and one for the Golgi, all based on the published RUSH method (Boncompain et al., 2012). We chose the Golgi-hook based on the original study as it did not obviously affect photoreceptor development or function. However, we found that expression of all four Streptavidin-hooks could affect the development of different tissues to a different degree, as discussed below. Next, we established a protocol for raising flies on Biotin-free food (addition of Biotin leads to release from the hook). Raising flies on Biotin-free food significantly affected offspring numbers, viability and caused significant developmental delays in particular during larval stages. Both effects of hook expression and Biotin-free food on development and viability limited the applicability of the RUSH system in our hands. In particular, we did not find conditions to study wing discs without adverse effects of the assay (Suppl. Fig. 5A-D). However, we identified conditions for the synchronized release method in photoreceptor neurons in *ex vivo* culture of developing fly brains (Ozel et al., 2015) as well as in salivary glands (Suppl. Fig. 5E-H).

We first performed proof-of-principle experiments using the well-characterized late endosomal Rab7 (Fig. 5B-E). The RUSH version of Rab7 (a fusion with Streptavidin binding protein (SBP)) rescued *rab7* mutant-associated lethality when expressed by the *rab7*-Gal4 knock-in, demonstrating that the SBP-fusion is a functional Rab GTPase. Rab7 forms transient rings marking the maturation of late endosomal compartments in photoreceptor cell bodies (Cherry et al., 2013). When raised on normal food, SBP-YFP-Rab7 localizes partly to the Golgi-hook and partly to characteristic endosomal rings (Fig. 5C). When raised on biotin-free food, almost all quantifiable YFP-Rab7 signal colocalized with the hook in both photoreceptors (Fig. 5B) and salivary glands (Suppl. Fig. 5E-H). Biotin incubation led to separation of SBP-YFP-Rab7 from the Golgi-hook within minutes and the formation of the characteristic Rab7-positive rings within 30-60 minutes (Fig. 5C-E). Based on the assay established for Rab7, we generated a complete collection of transgenic fly lines of Streptavidin-binding-protein (SBP)-tagged version for all 26 Rabs to facilitate comprehensive analyses and provide a generally available toolbox. We tested all 26 UAS-SBP-YFP-Rabs in comparison to the established UAS-YFP-Rab collection and exhibited identical subcellular localization patterns (Zhang et al., 2007) (nervous-system enriched Rabs shown in Suppl. Fig. 4).

Prior to Biotin addition, Rab7 forms aggregates together with the Golgi-hook, which resolve after release. Similar aggregates were observed for a subset of nervous system Rabs, in particular Rab3, Rab19, Rab26, Rab27, Rab32, RabX1 and RabX4 (Suppl. Fig. 6). Addition of Biotin leads to a reduction of these Golgi aggregates to varying degrees, followed by differential relocalization behavior within the first 15 minutes (Fig. 5F). Golgi aggregates had mostly dispersed after 60 minutes in the case of Rab3, Rab19 and Rab26, but significant aggregates remained for all others (Suppl. Fig. 5). 50% or more of the Golgi-hook compartments lost colocalization with RabX4, RabX1, RabX6 and Rab21 within 15 minutes of release (Fig. 5F,G). By contrast, only 20% or less of the hook compartments had released Rab19, Rab26 or Rab27 (Fig. 5F,H). Most of the ‘fast-releasing’ Rabs immediately marked distinct compartments in the cytoplasm (marked in green in Fig. 5F-H), whereas most ‘slow-releasing’ Rabs diffused in the cytoplasm (Fig. 5F). Three Rab proteins exhibited almost exclusive photoreceptor cell body membrane localization and could not be quantified in this way: Rab4, Rab9 and Rab23. Finally, Golgi-hook retention of Rab21 accumulated in large numbers of compartments in interommatidial cells and caused morphological irregularities that were not observed for any other Rab (Suppl. Fig. 6).

To test the extent to which the Rab proteins relocalized to axon terminals, we measured axon terminal/cell body fluorescence ratios (Fig. 5I-J). Significant increases of axon terminals localization were only observed for Rab4, Rab23 and Rab26 (Fig. 5J). However, in the case of Rab4 the ratio increase was exclusively due to the loss of hook localization, while Rab23 and Rab26 exhibited clear increases of the axon terminal signals (Fig. 5I). All three Rabs were either mostly membrane-bound (Rab4 and Rab23) or diffuse after their release (Rab26), providing no evidence for membrane compartment trafficking from the cell body to the axon terminals within 60 minutes. Notably, we observed low level Golgi-hook fluorescence and axon terminal localization of all Rabs to varying degrees already prior to synchronized release. The hook-bound axon terminal signal was highest for Rab3, Rab26, Rab27, Rab32 and RabX4. In sum, the RUSH acute release experiments identified ‘fast-releasing’ Rabs that are immediately recruited to membrane compartments in the cell body (RabX4, RabX1, Rab21), ‘slow-releasing’ Rabs that diffuse in the cytoplasm in the cell body (Rab26, Rab27, Rab19) and Rabs that are recruited to the axon terminal (Rab23 and Rab26).

### Integrative analysis of protein localization and mutant observations

In order to relate the localization of the nervous system Rab proteins to their possible roles during development and maintenance, we analyzed the localization of both endogenous and photoreceptor-expressed Rabs. First, we analyzed developing and adult optic lobes using a collection of endogenously tagged Rabs (Dunst et al., 2015). All 13 nervous system Rabs are expressed in varying patterns in the nervous system with predominant protein localization to synaptic neuropils (Suppl. Fig. 7), consistent with our previous analyses of tagged Rabs in the larval nervous system (Chan et al., 2011; Jin et al., 2012). In particular, widespread synaptic expression in adult neuropils is apparent for all Rabs except Rab27, Rab32, Rab23 and Rab9, which exhibit more cell-specific restrictions. The six mutants exhibiting stimulus-dependent functional maintenance defects all exhibit strong adult synaptic localization and are distinct from these four (Table 1C). These observations support the idea that the majority of Rabs with adult synaptic localization serve modulatory functions that become apparent under challenging conditions, namely Rab3, Rab26, Rab19, RabX6, Rab14, Rab30 and RabX4. By contrast, Rab27, Rab32, Rab23 and Rab9 are more likely to serve cell-specific functions, consistent with previous observations for each of the four in *Drosophila* (Chan et al., 2011; Dong et al., 2013; Gillingham et al., 2014; Ma et al., 2004)

Developmental localization is more varied and particularly strong in many cells for Rab3, Rab4, Rab19, Rab26, RabX1 and RabX4 (Suppl. Fig. 7). These include all three lines that exhibited developmental delays (RabX4, Rab19 and RabX1; Table 1D). By contrast, Rab21, Rab27, and Rab40 exhibit low expression in the pupal brain, consistent with normal developmental timing in the respective mutants described above. Rab32 appears specific to photoreceptor neurons; Rab9 and Rab23 appear highly restricted to a few cells in the adult brain. Hence, endogenous expression patterns of all nervous system-enriched Rabs suggest roles in different parts of the brain during development or function. Table 1 summarizes our findings from both functional and localization studies as described in more detail below.

In sum, our null mutant collection of all 26 Rab GTPases in *Drosophila* revealed that eight mutants are lethal or semi-lethal and two are infertile. Of the 18 viable null mutants (including the infertile ones), eight exhibit reduced viability under laboratory conditions. We found evidence for functional or maintenance defects based on larval progression or adult synaptic function for nine of the 18 viable Rabs (Table 1E). Six of these nine did not exhibit reduced viability under laboratory conditions and seven of these nine have previously been described as nervous-system enriched, and all of these localize to synapses. By contrast, we found evidence for developmental defects for only four of the 18, with three of the four exhibiting reduced viability. All four can be found in developing cell bodies, but have differential localization to developing and functional synapses (Table 1D).

The only Rabs for which our assays provided no evidence for a developmental or functional role are Rab9, Rab18, Rab21, and Rab39. These four also did not exhibit reduced viability. Of these only two have been previously characterized as neuronal-enriched: Rab9 exhibits mostly glia expression and Rab21 weak widespread expression in the adult brain.

All in all, we identified implications for 9 of the 13 nervous system-enriched Rabs during specific stages of developmental or functional maintenance. These defects are non-lethal under laboratory conditions, but represent sensitized genetic backgrounds that reveal limits to neuronal robustness under different challenging conditions.

### Loss of *rab26* does not discernibly affect membrane trafficking associated with synaptic vesicles or autophagy in the adult brain

Rab26 has been proposed to link synaptic vesicle recycling to autophagy based on experiments in mammalian cell culture and *Drosophila* (Binotti et al., 2015), which were based on the expression of GTP-locked and GDP-locked variants in the absence of a *rab26* null mutant. Our first mutant analysis presented here revealed that the *rab26* null mutant is viable. In support of the earlier hypothesis, we found that *rab26* is one of six *rab* null mutants that exhibit reduced stimulus-dependent functional maintenance (Fig. 3E); in addition, *rab26* null mutant axon terminals challenged with continuous light exhibited pronounced membrane accumulations (Fig. 4A). However, we found no significant changes of the autophagosomal marker Atg8/LC3 (Fig. 4B). These findings prompted us to probe in more detail putative roles of Rab26 at synaptic terminals.

First, we tested the GTP- and GDP-locked versions of Rab26 similar to experiments by Binotti et al. (2015). Expression of GTP-locked Rab26 in photoreceptor neurons leads to a complete loss of neurotransmission, while expression of GDP-locked or wild type Rab26, as well as the null mutant, do not block neurotransmission (Fig. 6A). Expression of wild type Rab26 marks distinct synaptic compartments that largely exclude synaptic markers (Syt1 and CSP; Fig. 6B-C) and the autophagosome marker Atg8 (Fig. 6D-E). By contrast, the recycling endosomal markers Rab11 (Fig. 6D-E) and the endosomal markers Hrs and Syx7 (Fig. 6F-G) all exhibit elevated levels. GTP-locked Rab26 protein forms more and enlarged accumulations at synaptic terminals as observed in the earlier study. These large Rab26-positive accumulations partly colocalize with early and recycling endosomal markers Syx7 and Rab11, but are again distinct from Atg8 or Syt1 labeling (Fig 6D-G). These findings suggest an endosomal role at synaptic terminals that may not be directly linked to synaptic vesicles and autophagy.

**Figure 6:**
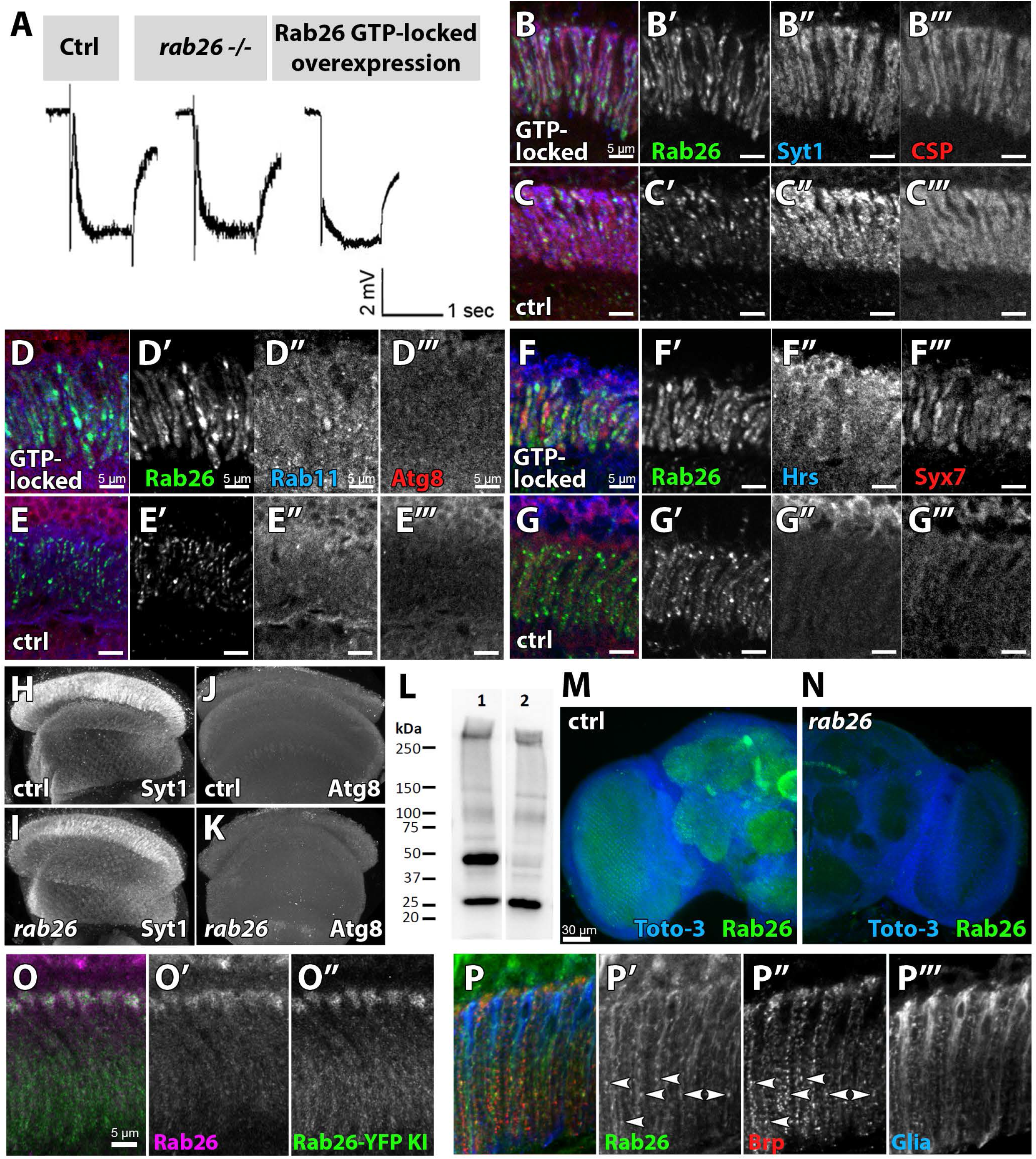
Loss of *rab26* does not discernibly affect membrane trafficking associated with synaptic vesicles or autophagy in the adult brain. (A) Representative ERG traces of recordings of newly hatched wild type, *rab26* mutant, and Rab26 GTP-locked overexpression flies. Only the Rab26 GTP-locked flies show a loss of ‘on’-transient. (B-G) Labeling of lamina cross-sections of Rab26 GTP-locked (B, D, and F) and control (C, E, and G) against Syt1 and CSP (B & C), Rab11 and Atg8 (D & E), and Hrs and Syx7/Avalanche (F & G). Scale bar = 5 μm; number of brains n = 3-5 per antibody staining. (H-K) Intensity comparison of optic lobes of newly hatched wild type and *rab26* mutant, stained against Syt1 (H & I) and Atg8 (J & K). Number of brains n = 3-5 per antibody staining. (L) Validation of the *rab26* null mutant by Western Blot with the newly generated Rab26 antibody. Wild type control shows the Rab26 band at around 45 kDa (1), which is lost in the *rab26* mutant (2). (M & N) Validation of the *rab26* null mutant by immunohistochemistry with the newly generated Rab26 antibody. The Rab26 antibody labels synaptic neuropil in different regions of wild type brains (green, M), which is lost in the *rab26* null mutant (N). Labeling of nuclei/ cell bodies with Toto-3 (blue). Scale bar = 30 μm; number of brains n = 3 per antibody staining. (O) Immunolabeling of Rab26 (red) shows high colocalization with the endogenously YFP-tagged Rab26 (green). Lamina cross-section of newly hatched flies. Scale bar = 5 μm; number of brains n = 3-5 per antibody staining. (P) Co-labeling of wild type lamina with Rab26 (green), Brp (synaptic marker, red) and ebony (glia marker, blue) reveals few synapses, positive for Rab26 and Brp in the proximal region of the lamina (white arrowheads, P’ and P”). No co-localization between Rab26 and ebony could be observed. Number of brains n = 3-5 per antibody staining.

Next, we compared the findings from GTP-locked Rab26 overexpression to the *rab26* null mutant. Adult brains mutant for *rab26* did not exhibit obvious changes of Atg8 or Syt1 (Fig. 6H-K). The null mutant brains appeared morphologically normal and exhibited no difference for any of the markers analyzed for GTP-locked Rab26 overexpression. These findings do not support a role in any essential membrane trafficking process during development and initial function.

To characterize neuronal Rab26-positive neurons and compartments in more detail, we generated a polyclonal antibody against the cytosolic N-terminus of Rab26 (see Materials and Methods). In Western Blots of whole brain homogenate, the Rab26 antibody labeled a 45 kDa band, consistent with predicted molecular weights between 41 kDa and 45 kDa, that is lost in the null mutant (Fig. 6L). Immunolabeling of brain tissue revealed labeling of synaptic neuropils at highly varying levels in different regions, which is lost in the null mutant (Fig. 6M-N) and colocalizes well with the endogenously tagged Rab26 (Fig. 6O). In the lamina, anti-Rab26 strongly labeled cell bodies distal of the photoreceptor axon terminals, and more weakly the axon terminal region (Fig. 6P). The distal lamina contains cell bodies of both glia and postsynaptic lamina neurons. Co-labeling with the glia marker ebony revealed no colocalization with Rab26. Instead, the synaptic marker Brp revealed a small subset of colocalizing synapses in the proximal regions of the axon terminals (Fig. 6P), i.e. the region of protein accumulations in our light-stimulation assay (Fig. 4A). Together, these observations raise the question whether the role of Rab26 is specific to a certain type of neuron or synapse.

### Rab26 is required for membrane receptor turnover associated with cholinergic synapses

Our *rab26* null mutant analyses have revealed a stimulus-dependent role in functional maintenance (Fig. 3E) associated with membrane protein accumulations at the proximal end of photoreceptor synaptic terminals (Fig. 4A). These mutant accumulations of the photoreceptor membrane protein Chaoptin became more pronounced with further increased (four days light) stimulation (Fig. 7A-B). To characterize the nature of these presynaptic protein accumulations, we tested a panel of 13 markers for membrane-associated proteins (Fig. 7A-M). We found the protein accumulations to be specifically enriched for the synaptic transmembrane cell adhesion molecule N-Cadherin (CadN) (Fig. 7C-D, M). By contrast, neither the autophagosomal marker Atg8, nor the synaptic vesicle marker Syt1 or the endosomal markers Rab5 and Rab7 were associated with the protein accumulations (Fig. 7I-L). Of the endosomal markers, only Syx7 was significantly increased (Fig. 7E, F, M). We conclude that continuous stimulation leads to the selective accumulation of presynaptic transmembrane receptors specifically in the most proximal part of photoreceptor terminals.

**Figure 7:**
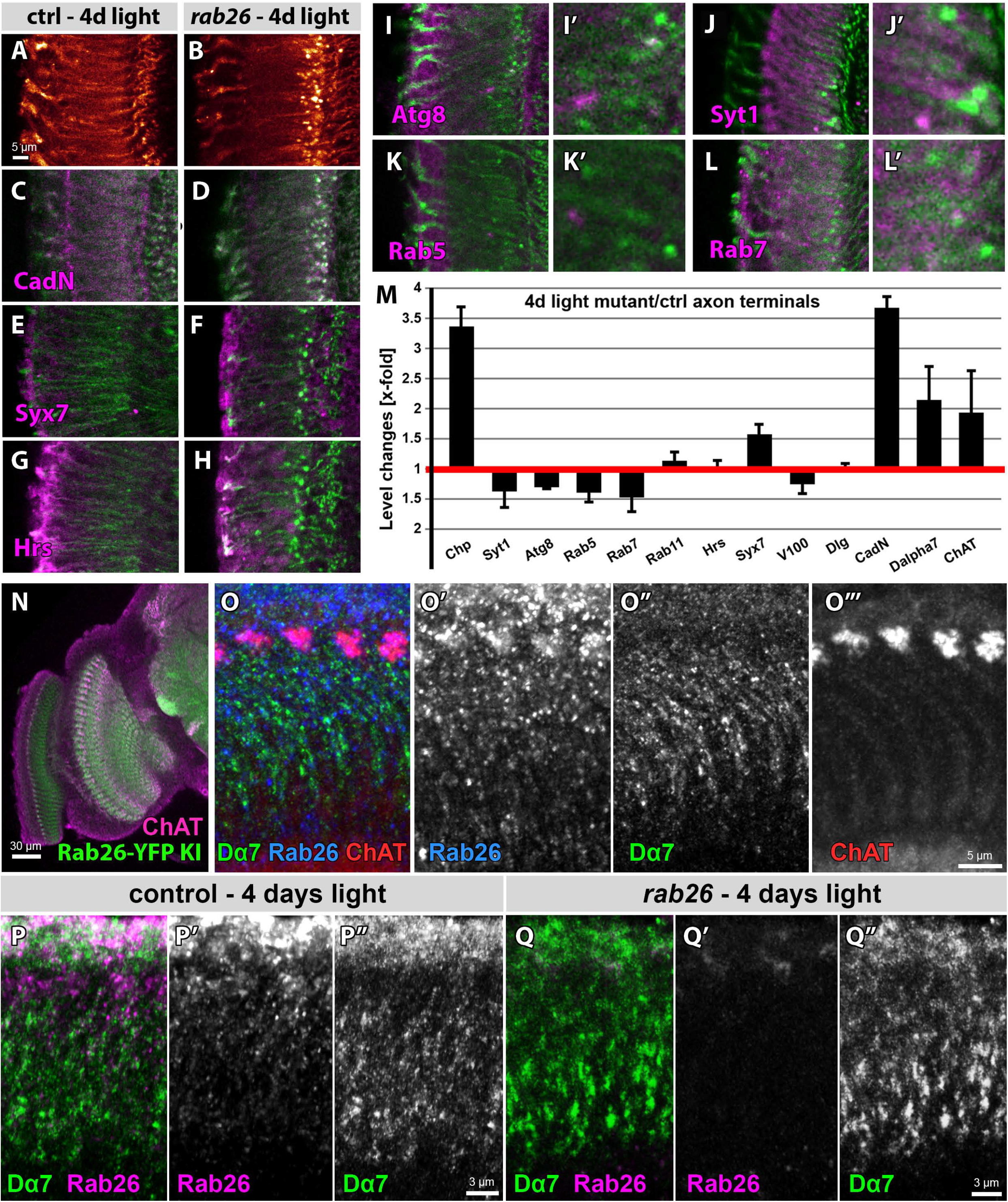
Rab26 is required for membrane receptor turnover associated with cholinergic synapses. (A-B) In contrast to the control (A), *rab26* mutant (B) shows Chaoptin-positive accumulations in the proximal lamina after 4 days light stimulation. Scale bar = 5 μm; number of brains n = 5 per antibody staining. (C-L) Chaoptin co-labeling (green) with CadN (C & D), Syx7/Avalanche (E & F), Hrs (G & H), Atg8 (I), and Syt1 (J), Rab5 (K), and Rab7 (L) in magenta. Only CadN is specifically enriched in the mutant accumulations (D), while levels of Syx7 are increased but show no co-localization with these accumulations (F). Shown are lamina cross-sections of wild type (C, E, and G) and *rab26* mutant (D, F, H, I-L) after 4 days light stimulation. Zoom-ins of Atg8- (I’), Syt1- (J’), Rab5- (K’), and Rab7-labeling (L’) reveal no enrichment in the Chaoptin accumulations. Number of brains n = 3-5 per antibody staining. (M) Quantification of level changes of 13 membrane-associated proteins in the *rab26* mutant axon terminals after 4 days light stimulation. Apart from Chaoptin (Chp), only CadN, Dα7 and ChAT are highly upregulated in the mutant axon terminals. Syx7 is upregulated as well but to a lesser extent. (N) Immunolabeling of newly hatched, endogenously YFP-tagged Rab26 with ChAT (magenta) reveals a similar pattern to Rab26 expression (green). Scale bar = 30 μm; number of brains n = 3-5 per antibody staining. (O) Co-labeling of newly hatched wild type lamina with Dα7 (green), Rab26 (blue) and ChAT (red). ChAT-positive L4 cell bodies are enriched for Rab26 (O’ & O’”), while Dα7-positive puncta in the lamina neuropil are often near but not colocalizing with Rab26 (O’ & O”). Scale bar = 5 μm; number of brains n = 3-5 per antibody staining. (P-Q) Co-labeling of wild type (P) and *rab26* mutant lamina (Q) with Dα7 (green) and Rab26 (magenta), shows Dα7 accumulations mostly in the proximal lamina after 4 days light stimulation. Scale bar = 3 μm; number of brains n = 3-5 per antibody staining.

Amongst lamina neurons, only L4 specifically forms synapses at the most proximal end of photoreceptor axon terminals (Fischbach and Dittrich, 1989; Luthy et al., 2014; Tadros et al., 2016). The function of L4 neurons remains unclear, but it was previously shown to express the vesicular acetylcholine transporter and choline acetyltansferase (ChAT) (Kolodziejczyk et al., 2008). We therefore tested whether Rab26-positive neurons are cholinergic using co-labeling of ChAT and the acetycholine receptor Dα7 (Fayyazuddin et al., 2006). Both proteins are expressed in a pattern very similar to Rab26 (Fig. 7N-O). In particular, ChAT labeling identified the L4 cell bodies, which are enriched for Rab26 (Fig. 7O). Anti-Dα7 receptor labeling revealed punctae in the lamina synaptic neuropil that are often near, but rarely directly colocalizing with Rab26 (Fig. 7O). After four days of light stimulation, both ChAT and Dα7 are significantly increased (Fig. 7M), with Dα7 accumulations most prominent in the proximal lamina (Fig. 7P-Q). In summary, our observations suggest that a requirement for Rab26 function in the first optic neuropil only becomes apparent after intense neuronal stimulation and is associated with cholinergic synapses. The observed Chaoptin, N-Cadherin and Dα7 receptor accumulations suggest a role in membrane receptor turn-over rather than synaptic vesicle recycling.

## DISCUSSION

In this study we generated a complete *rab* null mutant collection and focused on a functional analysis of those that are viable under laboratory conditions. Surprisingly, all previously described nervous system-enriched Rab GTPases fall into this category. However, challenging development with temperature or neuronal function with continuous stimulation revealed distinct requirement for most of these Rab GTPases. Our findings suggest that the majority of Rab GTPases modulate membrane trafficking in neurons and other tissues to maintain robust development and function under variable environmental conditions.

### A functional *rab* family profile

Rab GTPases have been analyzed in several comparative studies in order to gain a systematic view of membrane trafficking in cells (Best and Leptin, 2020; Chan et al., 2011; Dunst et al., 2015; Gillingham et al., 2014; Gurkan et al., 2005; Jin et al., 2012; Pfeffer, 1994; Stenmark, 2009; Zerial and McBride, 2001). All comparative studies to date have been based on expression profiling, the expression of GDP and GTP-locked Rabs or RNAi. However, we have previously described differences between loss of gene function and the expression of GDP-locked (often called dominant negative) variants (Chan et al., 2011; Cherry et al., 2013). The complete mutant collection will facilitate the comparison of molecularly defined null mutants with other functional perturbation approaches for all 26 Drosophila *rab* genes.

Our mutant analyses highlight that ‘viability’ is not a binary distinction of null mutants, but represents a continuous range of context-dependent phenotypes. Of the 26 null mutants, only 7 are fully lethal under any laboratory condition tested here *(rab1, rab2, rab5, rab6, rab7, rab8, rab11)*, while an eighth mutant is ‘semi-lethal’ based on few adult escapers *(rab35)*. Two more lines are viable, but infertile as homozygous adults *(rab10, rab30)*. Several others are highly sensitive to rearing conditions and may appear lethal depending on temperature, food, etc., including *rabX1, rabX4, rab19* and *rab32*. And another heterogeneous group of mutants exhibit reduced numbers of offspring or developmental or neuronal functional impairments depending on environmental conditions. Based on a non-quantitative survey of the literature, it seems likely that two-thirds or more of all *Drosophila* null mutants are viable under laboratory conditions, a ratio well reflected by *rab* mutants.

The eight (semi-)lethal mutants have all been previously analyzed in several model systems and contexts. Maybe not surprisingly, the majority of viable mutants contained all previously uncharacterized Rab GTPases. It is clear that the lethal mutants not analyzed here serve critical functions in development and function, including the nervous system. Our lists of development or functional Rabs (Table 1D, E) therefore only highlight non-essential roles under laboratory conditions. As such, these analyses provide a glimpse of the specialized roles under evolutionary selection that may represent functional specializations that correspond to environments typically encountered by *Drosophila*.

### Neuronal maintenance, membrane trafficking and the role of *rab26*

Our previous systematic analysis was based on expression profiling and suggested that the nervous system exhibits particularly pronounced expression of all Rab GTPases in *Drosophila* (Chan et al., 2011; Jin et al., 2012). We were surprised to find that all Rabs identified to be particularly enriched the nervous system proved to be viable under laboratory conditions. However, nervous system development and function has not only been evolutionarily selected for stereotypic precision, but also for flexibility and robustness to environmental challenges (Hiesinger and Hassan, 2018).

In this study we identified nine viable *rab* mutants affecting different aspects of neuronal function (Table 1E), and only four *rab* mutants affecting development (Table 1D). It is likely that key roles of Rab-dependent functions are executed by the lethal mutants not analyzed here. For example, *rab7* is a ubiquitously expressed gene, but disease-associated mutations primarily affect the nervous system and cause the neuropathy CMT2B (Cherry et al., 2013; Verhoeven et al., 2003). In axon terminals, local rab7-dependent degradation is required for turnover of membrane receptors, but not synaptic vesicles (Jin et al., 2018b). While null mutants for *rab7* are lethal, haploinsufficiency revealed neuronal sensitivity to reduced membrane degradation (Cherry et al., 2013). Similar to heterozygous *rab7*, our analyses of viable lines suggest that such evolutionarily selected functional properties may ‘hide’ in mutants that are characterized as viable under laboratory conditions.

Neurons require compartment-specific membrane trafficking in both axon terminals and dendrites (Jin et al., 2018a; Jin et al., 2018b). At presynaptic axon terminals, Rabs have been implicated in synaptic vesicle recycling, synaptic development and maintenance (Binotti et al., 2015; Graf et al., 2009; Sheehan et al., 2016; Uytterhoeven et al., 2011). We previously found that several neuron-enriched Rabs at axon terminals were positive for the recycling endosome marker Rab11 (Chan et al., 2011), including Rab26. Rab26 was subsequently identified as a possible link between autophagy and synaptic vesicle recycling (Binotti et al., 2015). Here we describe that *rab26* mutants exhibited neuronal functional defects only when challenged with continuous stimulation. However, we did not find obvious changes to autophagosomal and synaptic vesicle markers in the null mutant. Instead, Rab26 exhibited a highly specific expression pattern in the fly visual system with a particular enrichment in the L4 interneuron, which was previously shown to express cholinergic markers (Kolodziejczyk et al., 2008). While the function of L4 is unknown, it is not postsynaptic to photoreceptor neurons, but instead provides ‘feedback’ input on many cell types in the lamina (Luthy et al., 2014; Rivera-Alba et al., 2011; Tadros et al., 2016). Our finding of membrane receptor accumulations at the location of L4 synapses after continuous stimulation suggests a role that may depend on the intensity or duration of sensory input. Furthermore, the specific accumulations of the synaptic adhesion molecule N-Cadherin and the alpha7 nACh receptor suggest a role directly associated with cholinergic synapses in this system. Rab26 may thus represent the type of functionally relevant specialization necessary to endow the nervous system with robustness to variable and challenging environmental conditions (Hiesinger and Hassan, 2018). Hence, our study of *rab* mutants that are viable under laboratory conditions may help to elucidate an understanding of evolutionarily selected functional requirements under varying environmental conditions. The complete collection of null mutants and RUSH transgenic toolbox provide resources designed to facilitate such further studies.

## MATERIALS AND METHODS

### Fly husbandry and genetics

Flies were raised on molasses formulation food, with the exception of flies used in the RUSH-experiments. For RUSH experiments, flies were raised on egg white (avidin)-enriched molasses formulation food. The diet was modified by dissolving 20 g spray-dried, powdered egg white in 20 ml de-ionized water and added to 230 ml liquid food after the cooking process. The food temperature needs to be below 42°C to prevent coagulation of egg white. Stocks were kept on normal molasses formulation food at room temperature (22-23°C) in non-crowded conditions, which we defined as ‘normal laboratory conditions’. Flies were mostly raised at 25°C or 18°C and 29°C (developmental timing assay).

For the generation of CRISPR-targeted mutagenesis the following fly lines were used: w^1118^ and y^2^ cho^2^ v^1^; attP40{nos-Cas9}/CyO (NIG-FLY #CAS-0001). For the rescue of *rab30*^−^ infertility we used: *rab30^− Gal4-KI^*, UAS-YFP-Rab30WT.

For the analysis of the expression pattern of YFP- and SBP-YFP-tagged Rab wild type flies in P40% pupae the following *Drosophila* lines were used: YFP-tagged (GMR-Gal4/UAS-YFP-Rab3WT;GMR-myr-tdTomato/+, GMR-Gal4/UAS-YFP-Rab4WT;GMR-myr-tdTomato/+, GMR-Gal4/UAS-YFP-Rab9WT;GMR-myr-tdTomato/+, GMR-Gal4/UAS-YFP-Rab19WT;GMR-myr-tdTomato/+, GMR-Gal4/UAS-YFP-Rab21WT;GMR-myr-tdTomato/+, GMR-Gal4/UAS-YFP-Rab23WT;GMR-myr-tdTomato/+, GMR-Gal4/UAS-YFP-Rab26WT;GMR-myr-tdTomato/+, GMR-Gal4/UAS-YFP-Rab27WT;GMR-myr-tdTomato/+, GMR-Gal4/UAS-YFP-Rab32WT;GMR-myr-tdTomato/+, GMR-Gal4/UAS-YFP-Rab40WT;GMR-myr-tdTomato/+, GMR-Gal4/UAS-YFP-RabX1WT;GMR-myr-tdTomato/+, GMR-Gal4/UAS-YFP-RabX4WT;GMR-myr-tdTomato/+, and GMR-Gal4/UAS-YFP-RabX6WT;GMR-myr-tdTomato/+), SBP-YFP-tagged (GMR-Gal4/UAS-SBP-YFP-Rab3WT;GMR-myr-tdTomato/+, GMR-Gal4/UAS-SBP-YFP-Rab4WT;GMR-myr-tdTomato/+, GMR-Gal4/UAS-SBP-YFP-Rab9WT;GMR-myr-tdTomato/+, GMR-Gal4/UAS-SBP-YFP-Rab19WT;GMR-myr-tdTomato/+, GMR-Gal4/UAS-SBP-YFP-Rab21WT;GMR-myr-tdTomato/+, GMR-Gal4/UAS-SBP-YFP-Rab23WT;GMR-myr-tdTomato/+, GMR-Gal4/UAS-SBP-YFP-Rab26WT;GMR-myr-tdTomato/+, GMR-Gal4/UAS-SBP-YFP-Rab27WT;GMR-myr-tdTomato/+, GMR-Gal4/UAS-SBP-YFP-Rab32WT;GMR-myr-tdTomato/+, GMR-Gal4/UAS-SBP-YFP-Rab40WT;GMR-myr-tdTomato/+, GMR-Gal4/UAS-SBP-YFP-RabX1WT;GMR-myr-tdTomato/+, GMR-Gal4/UAS-SBP-YFP-RabX4WT;GMR-myr-tdTomato/+, and GMR-Gal4/UAS-SBP-YFP-RabX6WT;GMR-myr-tdTomato/+).

For the proof-of-principle experiments of the RUSH system the following *Drosophila* line was used: (GMR-Gal4/UAS-SBP-YFP-Rab7;UAS-Streptavidin-Golgin84/+). For the biotin-release RUSH-experiments the following *Drosophila* lines were used: (GMR-Gal4/UAS-SBP-YFP-Rab3;UAS-Streptavidin-Golgin84/+, GMR-Gal4/UAS-SBP-YFP-Rab4;UAS-Streptavidin-Golgin84/+, GMR-Gal4/UAS-SBP-YFP-Rab9;UAS-Streptavidin-Golgin84/+, GMR-Gal4/UAS-SBP-YFP-Rab19;UAS-Streptavidin-Golgin84/+, GMR-Gal4/UAS-SBP-YFP-Rab21;UAS-Streptavidin-Golgin84/+, GMR-Gal4/UAS-SBP-YFP-Rab23;UAS-Streptavidin-Golgin84/+, GMR-Gal4/UAS-SBP-YFP-Rab26;UAS-Streptavidin-Golgin84/+, GMR-Gal4/UAS-SBP-YFP-Rab27;UAS-Streptavidin-Golgin84/+, GMR-Gal4/UAS-SBP-YFP-Rab32;UAS-Streptavidin-Golgin84/+, GMR-Gal4/UAS-SBP-YFP-Rab40;UAS-Streptavidin-Golgin84/+, GMR-Gal4/UAS-SBP-YFP-RabX1;UAS-Streptavidin-Golgin84/+, GMR-Gal4/UAS-SBP-YFP-RabX4;UAS-Streptavidin-Golgin84/+, and GMR-Gal4/UAS-SBP-YFP-RabX6;UAS-Streptavidin-Golgin84/+). For the RUSH-release experiments in imaginal wing disc pouch and 3^rd^ instar salivary glands, the following lines were used: UAS-SBP-YFP-Rab7/nubbin-Gal4; UAS-Streptavidin-Golgin84/+ and UAS-SBP-YFP-Rab7/+; UAS-Streptavidin-Golgin84flies/sgs3-Gal4.

For the analysis of the expression pattern of endogenously tagged Rab GTPases in pupae and one day old adults, the following homozygous *Drosophila* lines were used: (EYFP-Rab3, EYFP-Rab4, EYFP-Rab9, EYFP-Rab19, EYFP-Rab21, EYFP-Rab23, EYFP-Rab26, EYFP-Rab27, EYFP-Rab32, EYFP-Rab40, EYFP-RabX1, EYFP-RabX4 (EYFP-RabX4/TM6B for adult brain analysis), and EYFP-RabX6, Dunst et al., 2015).

For the analysis of the identity of the Chaoptin-positive accumulations in rab26^−^ lamina after 4 days of light stimulation, following *Drosophila* lines were used:;;rab26^−^ and yw as wild type control. For the analysis of possible co-localization between Rab26-positive compartments and synaptic vesicle markers as well as endomembrane trafficking markers, following *Drosophila* lines were used:;sGMR-Gal4/UAS-YFP-rab26WT; and;sGMR-Gal4/UAS-YFP-Rab26CA;. For the comparison of the anti-Rab26 antibody labeling with the YFP-knock in line, the following *Drosophila* line was used:;UAS-YFP-Rab26WT/+; rab26^exon1^-Ga14/+.

### Generation of null mutant flies

All CRISPR/Cas9-mediated *rab* mutants, except *rab18*^−^ and *rab26*^−^, were generated by WellGenetics Inc. (Taipei, Taiwan), by homology-dependent repair (HDR) using two guide RNAs and a dsDNA plasmid donor (Kondo and Ueda, 2013). Briefly, upstream and downstream gRNA sequences were cloned into a U6 promoter plasmid. For repair, a cassette, containing two loxP-sites flanking a 3xP3-RFP with two homology arms was cloned into a donor template (pUC57-Kan). A wild type strain (w^1118^) was injected with the donor template as well as specific *rab*-targeting gRNAs and hs-Cas9. F1 progeny positive for the positive selection marker, 3xP3-RFP, were further validated by genomic PCR and sequencing. The CRISPR null mutants were validated as described in the next section. gRNA sequences as well as specifics on the different CRISPR mutants are as follows:

***rab4***^−^: Replacement of 944 bp region, −100 bp relative to ATG to +4 bp relative to the first bp of *rab4* stop codon, by floxable cassette. This results in the deletion of the entire coding region and part of 5’-UTR.Upstream gRNA sequence: AATCCGATAGTCCTGAAGTC, downstream gRNA sequence: TAAACGCGACAGGTGCAATC.
***rab9***^−^: Replacement of 2446 bp region, +98 bp relative to ATG to +111 bp relative to the first bp of *rab9* stop codon, by floxable cassette. Upstream gRNA sequence: GTTGTTCTCCTCGTAGCGAT, downstream gRNA sequence: ATTCCAGTCCGCGGAGGGGC.
***rab10***^−^: Replacement of 1644 bp region, +57 bp relative to ATG to +70 bp relative to the fist bp of *rab10* stop codon, by cassette, which contains 3 stop codons upstream of floxable 3xP3-RFP. Upstream gRNA sequence: CTGATCGGTGATTCAGGAGT, downstream gRNA sequence: GAACGGGGCGTGGTTTGGCC.
***rab14***^−^: Replacement of 930 bp region, −17 bp relative to ATG of *rab14-RB* isoform to −61 bp relative to the first bp of *rab14* stop codon, by floxable cassette. Upstream gRNA sequence: GATGAGCAAAGTGCGCAGCG, downstream gRNA sequence: GAAGTTCGCGACGGCTGCGA.
***rab21***^−^: Replacement of 608 bp region, +12 bp relative to ATG of *rab21-RD* isoform to −109 bp relative to first bp of *rab21* stop codon, by floxable cassette. Upstream gRNA sequence: CAATGAGCTCGAGCAGAACG, downstream gRNA sequence: GACTCGCATCCGGTTGCCGT.
***rab23***^−^: Replacement of 1700 bp region, −35bp relative to ATG to +173bp relative to the first bp of *rab23* stop codon, by floxable cassette. Upstream gRNA sequence: CAATCAAACACCTGGGCGAG, downstream gRNA sequence: CATGTCTGAACCACATCACG.
***rab35***^−^: Replacement of 816 bp region, −24 bp relative to ATG of *rab35-RC* isoform to +20 bp relative to the first bp of *rab35* stop codon, by floxable cassette. Upstream gRNA sequence: CAGCAATGTCATATGCCGAA, downstream gRNA sequence: AGGTGAAAGCGGCTCCGGCA.
***rab39***^−^: Replacement of 898 bp region, +92 bp relative to ATG to −93 bp relative to the first bp of *rab39* stop codon, by floxable cassette. Upstream gRNA sequence: CACAGACGGCAAATTCGCCG, downstream gRNA sequence: TCGATCCGGCGAATATAAGG.
***rab40***^−^: Replacement of 1407 bp region, +2 bp relative to ATG to −93 bp to the first bp of *rab40* stop codon, by floxable cassette. Upstream gRNA sequence: CCTTGGTCATGGTTCCCATG, downstream gRNA sequence: TTGAGCGTCGACTTCACCGA.
***rabX4***^−^: Replacement of 962 bp, −2 bp relative to ATG to −61 bp to first bp of *rabx4* stop codon, by floxable cassette. This results in the deletion of the entire coding sequence. Upstream gRNA sequence: CTCCGCCAGCTCCGTCAACA, downstream gRNA sequence: AAGAAATCACCCGGCTCCAA.
***rab18***^−^: For the generation of the *rab18* null mutant, first a *rab18* sgRNA-expressing plasmid (pBFv-U6.2-rab18-sgRNA) was generated. For this, *rab18* sgRNA sequence 5’-GGTGATCGGGGAAAGCGGCG (directly after the *rab18* start codon) was cloned into BbsI-digested pBFv-U6.2 plasmid. Second, a pCR8-rab18-3xP3-RFP plasmid was generated by soeing PCR and restriction enzyme digestion. For this, two 500 bp homology arms (HA) around the *rab18* sgRNA targeting site were amplified, using the following primers: left HA fwd: TCCTAAATTTATGATATTTTATAATTATTT; left HA rev: CTGGACTTGCCTCGAGTTTTTTAGATCTGTGTGGTTTGAGCTCCGCTT; right HA fwd: CAAACCACACAGATCTAAAAAACTCGAGGCAAGTCCAGGTGCAGTCCC; right HA rev: CGAACTGATCGCATTTGGCT. The resulting PCR product was then cloned into pCR8 vector (pCR8-rab18LA+RA). The 3xP3-RFP cassette, containing 3 stop codons upstream of the RFP, was cloned into pCR8-rab18LA+RA by *Bgl*II and *Xho*I double digestion to get the final pCR8-rab18-3xP3RFP plasmid. *Nanos-Cas9* fly embryos were co-injected with the two plasmids pBFv-U6.2-rab18-sgRNA and pCR8-rab18-3xP3RFP. F1 progeny positive for the selection marker, 3xP3-RFP, were further validated by genomic PCR.
***rab26***^−^: Replacement of 9760 bp region, −125 bp relative to ATG to +1310 bp to the end of coding exon 2, by positive selection marker 3xP3-dsRed flanked by loxP-sites. This leads to the complete deletion of ATG1 (exon 1) and ATG2 (exon 2) of *rab26* gene. Briefly, a *rab26* sgRNA-expressing plasmid was generated by cloning the *rab26* sgRNA 5’-GACAGTTTCGGAGTTAATTA into a Bbsl-digested U6-BbsI-chiRNA plasmid (addgene, plasmid #45946, donated by Kate O’Connor-Giles lab). *Nanos-Cas9* fly embryos were co-injected with the *rab26* sgRNA containing U6-chiRNA plasmid and the pHD-DsRed-attP plasmid (donated by Kate O’Connor-Giles lab). F1 progeny positive for the selection marker, 3xP3-dsRed, were further validated by genomic PCR.

In addition, six new *rab* mutants were generated by ends-out homologous recombination based on previously generated Gal4 knock-ins in large genomic fragments (Chan et al., 2011). All *rab* mutants generated by ends-out homologous recombination are ‘ORF knock-ins’ (replacing the entire open reading frame), except for rab4^−^, which is an ‘ATG knock-in’ (replacing the first exon including the start codon). The methods used for the replacements in the endogenous loci have been described previously in detail (Chan et al., 2012; Chan et al., 2011).

### Verification of *rab* null mutants by PCR

The newly generated *rab* null mutants were confirmed by genomic PCR, either using Phusion High-Fidelity PCR Kit (Thermo Scientific) (majority of rab mutants) or the SapphireAmp^®^ Fast PCR Master Mix (TaKaRa) (*rab26*)^−^. The following primer pairs, flanking the gene or inserted cassette, were used for the validation: *rab2*^−^ (Fwd: 5’-TGGCCACACTGTCGCTAGCC and Rev: 5’-CGCCTCCTCTACGTTGGCAG), *rab4*^−^ (Fwd: 5’-GGTTTTGATCGTGTCCTGCG and Rev: 5’-AGACAACTCTTACCGCTGCC), *rab9*^−^ (Fwd: 5’-GGCACTATGACGAACATGCGG and Rev: 5’-tttgcagcactgggaaatccg), *rab10*^−^ (Fwd: 5’-atatctcttgtcacctgcgcc and Rev: 5’-cgaccaccatccatcgttcgg), *rab18*^−^ (Fwd: 5’-AAACAAAGCAGCAAGGTGGC and Rev: 5’-CTCCTCGTCGATCTTGTTGCC), *rab19*^−^ (Fwd: 5’-CCAGTTAACGGCCAGAACGG and Rev: 5’-TTGCCTCTCTGAGCATTGCC), *rab21*^−^ (Fwd: 5’-CAATGGGAACGGCTAAATGCC and Rev: 5’-caacatttaTCGCCGAGTGCC), *rab23*^−^ (Fwd: 5’-CACCTGCCGGCTTAGATGCG and Rev: 5’-GAGATATCGGAACCGGCCCG), *rab26*^−^ (Fwd: 5’-CGATGAAGTGGACATGCACCC and Rev: 5’-tgcacttgaacttcactggcg), *rab30*^−^ (Fwd: 5’-ACCCAGCGACTCAAAAACCC and Rev: 5’-GCTGCACAGTTTCCAGATCCG), *rab35*^−^ (Fwd: 5’-CGAATCGTAAGCCAAGAACCC and Rev: 5’-ACTAATGGTGACGCACTGGC), *rab39*^−^ (Fwd: 5’-TAACAACCACCAGCGACAGCC and Rev: 5’-CGTATACCTCGTGTGACTGGC), *rab40*^−^ (Fwd: 5’-caatgagtaaacccctagcgg and Rev: 5’-TGGGTATGGGTATGGTATGGG), *rabX1*^−^ (Fwd: 5’-GTGCCCAAGAAATCAGACGC and Rev: 5’-AGTCAGATGGGCTTAGAGCG), *rabX4*^−^ (Fwd: 5’-CTGTAACCGAAAACCTCCGC and Rev: 5’-CAACTTGCTCAGGTTCTGCG), and *rabX6*^−^ (Fwd: 5’-GTCGCACTGTTGTTGTCGCC and Rev: 5’-CTCTGCGTGAGCATTGAGCC). For the validation of the mutants generated by homologous recombination the following cassette specific primers were used: Reverse primer in Gal4-region: 5’-CGGTGAGTGCACGATAGGGC (*rab2*^−^, *rab4*^−^, *rabX1*^−^), second reverse primer in Gal4-region: 5’-CAATGGCACAGGTGAAGGCC (*rab19*^−^, *rab30*^−^, *rabX6*^−^). The following cassette specific primers were used for the validation of CRISPR-generated null mutants: Reverse primer in RFP-region: 5’-GCTGCACAGGCTTCTTTGCC (*rab9*^−^, *rab10*^−^, *rab18*^−^, *rab39*^−^, *rabX4*^−^), second reverse primer in RFP-region: 5’-ACAATCGCATGCTTGACGGC (*rab21*^−^, *rab35*^−^, *rab40*^−^), forward primer in RFP-region: 5’-GGCTCTGAAGCTGAAAGACGG *(rab23)*^−^, forward primer in dsRed-region: 5’-ATGGTTACAAATAAAGCAATAGCATC (*rab26*^−^) and reverse primer behind right-arm of inserted dsRed-cassette: 5’-AAACCACAGCCCATAGACG.

The CRISPR null mutants were independently validated in our lab and by WellGenetics Inc. (Taipei, Taiwan).

### Generation of the RUSH toolbox

All RUSH-flies were generated by WellGenetics Inc. (Taipei, Taiwan), by conventional cloning and phiC31 integrase-mediated transgenesis. For the generation of the SBP-YFP-tagged Rab GTPases, two separate fragments were amplified by PCR, YFP-rab fragment from the genomic DNA of the respective UAS-YFP-RabWT lines (Zhang et al., 2007) and SBP fragment (addgene, plasmid #65305, donated by Franck Perez). Both fragments were cloned into the pUAST-attB vector, using Xhol/Xbal sites and the constructs were integrated into the same landing site y^1^w^1118^; PBac{y^+^-attP-3B}VK00002 (Bloomington stock #9723) in the *Drosophila* genome. SBP-YFP-tagged lines were generated for all 26 *Drosophila* Rab GTPases. For the generation of the Streptavidin-tagged hook line, the Streptavidin-Golgin84 fragment (addgene, plasmid #65305, donated by Franck Perez) was amplified by PCR and cloned into the pUAST-attB vector, using Xhol/Xbal sites. Two different landing sites were used for the integration of the construct into the *Drosophila* genome, second (y^1^w^1118^; PBac{y^+^-attP-9A}VK00018 (Bloomington stock #9736)) and third chromosome (y^1^ M{vas-int.Dm}ZH-2A w*; PBac{y^+^-attP-3B}VK00033 (Bloomington stock #24871)).

Three other hooks, namely Ii-, KDEL-, and STIM1-NN-hook, were generated as well, all located at the Endoplasmic Reticulum (not used in this study). The same method as for the generation of the Golgi-hook and the same two landing sites were used. These plasmids were used here: Streptavidin-Ii fragment (addgene, plasmid #65312), Streptavidin-KDEL (addgene, plasmid #65306), and Streptavidin-STIM1-NN (addgene, plasmid # 65311), and all were donated by the Frank Perez lab. Golgin84- as well as Ii-hook are facing the cytoplasmic site, while KDEL- and STIM1-NN-hook are facing in the luminal direction.

### Immunohistochemistry

Pupal and adult eye-brain complexes were dissected and collected in ice-cold PBS. The tissues were fixed in PBS with 4% paraformaldehyde for 30 minutes and washed in PBST (PBS + 1% Triton X-100). The following primary antibodies were used: rabbit anti-Rab5 (1:1000, Abcam), rabbit anti-Rab7 (1:1000, gift from P. Dolph), mouse anti-Rab11 (1:500, BD Biosciences), guinea pig anti-Rab26 (1:2000, made for this study), rabbit anti-Syt1 (1:1000, DSHB), rabbit anti-GABARAP+GABARAPL1+GABARAPL2 (ATG8) (1:100, Abcam), rabbit anti-Syx7/Avalanche (1:1000, gift from H. Krämer), guinea pig anti-Hrs (1:300, gift from H. Bellen), rabbit anti-DPAK (1:2000) mouse anti-DLG (1:100, DSHB), rat anti-Dα7 (gift from H. Bellen), rat anti-nCadherin (1:100, DSHB), guinea pig anti-v100 (1:1000, Hiesinger lab), mouse anti-CSP (1:50, DSHB), mouse anti-ChAT (1:100, DSHB), mouse anti-nc82 (1:20, DSHB), rabbit anti-ebony (1:200) and mouse anti-Chaoptin (1:50, DSHB). Secondary antibodies used were Donkey anti-mouse Alexa 405, Goat anti-mouse Alexa 488, Goat anti-guinea pig Alexa 488, Goat anti-rabbit Alexa 647, Goat anti-guinea pig Cy5 (1:500; Jackson ImmunoResearch Laboratories), and Toto-3 (1:1000; Thermo Fisher) as nuclear counterstain.

For RUSH-experiments, pupal eye-brain complexes were dissected in ice-cold Schneider’s *Drosophila* Medium and collected in ice-cold culture medium (Ozel et al., 2015). Experimental tissue was incubated in culture medium containing D-Biotin (working solution 2 mg D-Biotin/1 ml culture medium) for either 15 or 60 minutes (15, 30, 60 and 120 minutes for the proof-of-principle experiments; 15 and 30 minutes for release experiments in salivary glands). The control and experimental eye-brain complexes were washed with Schneider’s *Drosophila* Medium after biotin-incubation, fixed in PBS with 4% paraformaldehyde for 30 minutes and washed with PBST (PBS + 0.4% Triton X-100). The following primary antibody was used: mouse anti-Streptavidin (1:100, Novus Biologicals) in combination with the following secondary antibodies: Goat anti-mouse Alexa 647 (1:500; Jackson ImmunoResearch Laboratories) or Goat anti-rabbit Alexa 594 (1:800, Thermo Fisher). Salivary glands were incubated in the primary antibody solution for three days, while incubation time with all other antibodies was overnight. DNA was visualized using Hoechst staining dye (1:2000, Thermo Fisher).

All samples were mounted in Vectashield mounting medium (Vector Laboratories). To fully expose lamina photoreceptor terminals, pupal brains were mounted with their dorsal side up.

### Generation of rab26 antibody

The cDNA sequence corresponding to amino acids 1-192 of *rab26* was amplified by PCR and cloned into the pET28a (Invitrogen) vector for protein expression. Guinea pig antibodies against this domain were raised by Cocalico Biomedicals, Inc. using the purified recombinant protein.

### Confocal Microscopy, Image Processing and Quantification

All microscopy was performed using a Leica TCS SP8 X (white laser) with 20x and 63x Glycerol objectives (NA = 1.3). Leica image files were visualized and processed using Imaris (Bitplane) and Amira (Thermo Fisher). Postprocessing was performed using ImageJ (National Institute of Health), and Photoshop (CS6, Adobe, Inc).

Quantification of relative signal levels (RUSH cell body/axon terminals; axon terminal antibody labeling with and without light stimulation) was performed by measuring 8 bit pixel intensities in multiple regions of interest, followed by statistical analyses.

For co-localization experiments, all quantification was performed manually on single slices and only individually discernible compartments were counted. The statistical analyses were performed using RStudio (RStudio Inc.) and GraphPad Prism 8.3.0 (GraphPad Software, Inc.), and the specific statistical tests used as well as sample numbers for experiments are indicated in the respective figure legends.

### Biochemistry

Proteins were extracted from 20 adult fly brains per genotype in RIPA buffer containing 150 mM NaCl, 0.1% Triton X-100 (Sigma), 0.1% SDS (Amresco), 50 mM Tris-HCL and 1x complete protease inhibitors (Sigma), pH 8. Samples were incubated on ice for 20 minutes and centrifuged at 16 000 RCF, 10 minutes at 4°C to remove cell debris. Laemmli buffer (Biorad) was added to the supernatant. After incubation for 5 minutes at 95°C, the samples were loaded on a 4-15% SDS-polyacrylamide gel (Biorad) and then transferred to PVDF membrane (Biorad). Primary antibody used was rabbit anti-Rab26 (1:1000) and corresponding secondary was used 1:5000. The signals were detected with Clarity Western ECL (Biorad).

### Developmental assays

For the analysis of developmental timing of homozygous, viable *rab* mutants, three crosses with equal number of flies (ratio female to male ~2:1) and same genotype were set up a few days prior to the start of the experiment, to ensure good egg laying. Of each of those, again three equal groups were formed and egg laying was allowed for 24 hours at room temperature. Egg containing vials were then shifted to the respective temperatures (18°C, 25°C, or 29°C), while the parental flies remained at room temperature for the duration of the experiment. The shifting of egg containing vials was repeated 6 more times, leading to a total of 21 ‘experimental’ vials per temperature per genotype. Developing flies were kept at the respective temperatures until three days after they hatched, and the total number of hatched offspring was counted.

To study the effect of temperature stress on fly wing development, *rab* null mutants were reared at 18°C and 29°C. All mutant lines were set up with 10 females and 3 males and kept in their vials for 48 hours of egg laying, so as to prevent overcrowding in the vials. Adult female flies were collected not earlier than 24 hours after eclosion and placed in a 1:1 solution of glycerol:ethanol for a minimum of several hours, after which the wings were removed at the joint and mounted in the same solution. Wings were imaged with Zeiss Cell Observer microscope and their size measured in FIJI.

### Neuronal stimulation and electroretinogram (ERG) recordings

Newly eclosed adults were either placed in a box for constant white light stimulation or placed in light-sealed vials (in the same box) for constant darkness. The lightbox contains two opposing high-intensity warm white light LED-stripe panels, each emitting ~1600 lumen (beam angle = 120°, distance between light source and vials = 16 cm). Temperature (22°C) and humidity (59%) inside the box were kept constant. Flies were kept inside the box for up to 7 days (wild type sensitization curve) or for 2 and 4 days (function and maintenance experiments).

For the ERG recordings, the flies were anesthetized and reversibly glued on microscope slides using non-toxic school glue. The recording and reference electrodes were filled with 2 M NaCl and placed on the retina and inside the thorax. Flies were exposed to a series of 1 second light/dark pulses provided by a computer-controlled white light-emitting diode system (MC1500; Schott) as previously reported (Cherry et al., 2013). Two different light stimulus intensities, dim (5.29e^13^ photons/cm^2^/sec) and bright (1.31e^16^ photons/cm^2^/sec), were used. Retinal responses were amplified by a Digidata^®^ 1440A, filtered through a Warner IE-210, and recorded using AxoScope 10.6 by Molecular Devices. All ERG recordings were performed in non-pigmented, white-eyed flies, which are more sensitive to light stimulation than pigmented ones. 25-30 flies were examined for each genotype, condition, and time point.

## Acknowledgements

We would like to thank members of the Boutros, Hiesinger, Wernet and Hassan labs for their support and helpful discussions. We thank Hugo Bellen, Helmut Krämer and the Developmental Hybridoma Bank for reagents and Gerit Linneweber for comments on the manuscript. This work was supported by grants from the NIH (RO1EY018884), the German Research Foundation (DFG, SFB/TRR186) to P.R.H. and the German Research Foundation (DFG, SFB/TRR186) to M.B.

## Supplemental Figure Legends

**Suppl. Figure 1: Design of newly generated *rab* mutants.**

(A and C) Schematic depiction of the inserted knock-in cassettes. For ends-out homologous recombination a Gal4-3xP3-RFP-Kanamycin cassette, with loxP-sites flanking the 3xP3-RFP-Kan region, was inserted. For CRISPR/Cas9-mediated mutagenesis a 3xP3-RFP- or 3xP3-dsRed (for *rab26)* cassette, flanked by loxP-sites, was inserted.

(B and D) Schematics of genomic loci as depicted on FlyBase GBrowse (https://flybase.org/cgi-bin/gbrowse2/dmel/). The exon/intron region, with exon as wide orange bars, introns as black lines and 5’ UTRs and 3’UTRS as grey wide bars. The red half-arrows highlight regions replaced for ‘ORF knock-ins’ (B) or ‘CRISPR knock-ins’ (D); blue half-arrows highlight regions replaced for ‘ATG knock-ins’ *(rab4* in B).

**Suppl. Figure 2: Examples of wing defects after development at different temperatures.**

(A-H) Wing sizes of *rab* mutants at 18°C and 29°C. Flies at 29°C have on average 30% smaller wings than flies at 18°C (A-B). At 18°C, *rabX1* has significantly larger wings than control, while *rab19* has significantly smaller wings than control (C, E). At 29°C, rab9 has larger wings than control, while *rabX6* has smaller wings than control (D, F). *Rab23* shows, in addition to the PCP phenotype that is consistent at both temperatures, a p-cv vein shortening that is present in 90% of cases at 18°C (G, I), but is reduced to 12% at 29°C (H, J). Scale bar = 500 μm (A-H), 100 μm (I-J).

**Suppl. Figure 3: Systematic analysis of photoreceptor axon morphology of newly eclosed adults and after 2 days continuous light stimulation.**

(A) Labeling of newly hatched wild type and mutant photoreceptor projections with Chaoptin reveals no noticeable morphological differences. Chaoptin-positive accumulations in non-photoreceptor cells are visible in *rabX1*. Optic lobe overview (top panel), lamina cross-section with R1-R6 axon terminals (middle panel), and R7-R8 axon terminals (bottom panel). Scale bar top panel = 20 μm, middle and bottom panel = 5 μm; number of brains n = 3-5 per antibody staining.

(B) Labeling of wild type and mutant photoreceptor projections with Chaoptin after 2 days light stimulation. Chaoptin-positive accumulations in non-photoreceptor cells are visible in *rabX1*. Only *rab19* and *rab26* display morphological differences in their photoreceptor projection terminals, showing membrane accumulations in the tips of R1-R6 axon terminals. Optic lobe overview (top panel), lamina cross-section with R1-R6 axon terminals (middle panel), and R7-R8 axon terminals (bottom panel). Scale bar top panel = 20 μm, middle and bottom panel = 5 μm; number of brains n = 3-5 per antibody staining.

**Suppl. Figure 4: Comparison of newly generated Rab-YFP-SPB fusion proteins to Rab-YFP fusion proteins when expressed in photoreceptor neurons.**

Expression of YFP-tagged and SBP-YFP-tagged Rabs (green) shows no major differences in their subcellular localization patterns in ~P+40% pupal brains. Inverted channel shows expression of YFP- and SBP-YFP-tag. Labeling of photoreceptor projections by GMR-myr-tdTomato (red). Scale bar = 20 μm; number of brains n=3-6.

**Suppl. Figure 5: RUSH-associated defects in the wing disc and proof-of-principle experiment in salivary glands.**

(A-D) RUSH expression in wing imaginal discs. Larvae expressing the Golgi-hook (A) or the RUSH-Rab7 (B) under the nubbin-Gal4 driver show healthy organs on normal fly food. Dual overexpression of these constructs on normal fly food shows apoptosis and dead cell shedding, with wings of adult flies possessing notches and other deformities (C). Overexpression on biotin-deficient fly food results in highly deformed wing discs and larval death before pupation (D). Scale bar = 50 μm. (E-H) Expression of Golgi-hook and RUSH-Rab7 in salivary glands under the sgs3-Gal4 driver resulted in healthy organs when flies were reared on both normal (E) and biotin-deficient food (F). Release of RUSH-Rab7 from the Golgi-hook was achieved by biotin-incubation for 15 (G) and 30 minutes (H). Scale bar = 50 μm

**Suppl. Figure 6: RUSH release experiments in all nervous-system enriched *rab* mutants.**

Retention and release of ‘nervous system high’ Rabs from the Golgi-hook in ~P+40% pupal photoreceptor cell bodies after 60 minutes biotin-incubation. Scale bar = 2 μm; number of retinae n = 3-5 for each incubation time point.

**Suppl. Figure 7: Expression patterns of nervous-system enriched Rabs based on endogenously tagged Rabs generated by Dunst el al., 2015.**

(A-B) Expression pattern of EYFP-tagged Rabs (green) in ~P+40% pupal brains (A) and newly hatched adult brains (B). Immunolabeling of pupal photoreceptor projections with Chaoptin (red). Inverted channel shows expression of EYFP-tag. Scale bar = 20 μm (A), 30 μm (B); number of brains n = 3-6 per developmental stage.

**Suppl. Table 1: Quantitative analysis of the developmental timing assay at different temperatures.**

Summary of developmental time for wild type and all fertile, homozygous viable *rab* mutants at 18°C, 25°C and 29°C. Listed are number of days (after 24 hours of egg collection) until first 1st instar larvae, pupae or adults appear, as well as total number of adults hatched and number of adults per vial. Days are given in mean ± SEM.

