## Supplementary Figures and Table for "Systematic functional analysis of Rab GTPases reveals limits of neuronal robustness in *Drosophila*"

### Supplementary Figure 1

A Homologous recombination knock-in cassette

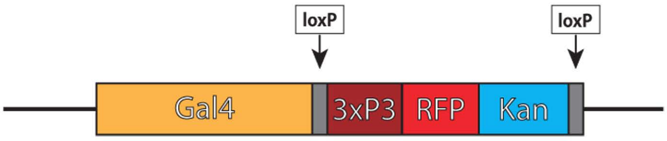


B Homologous recombination mutants

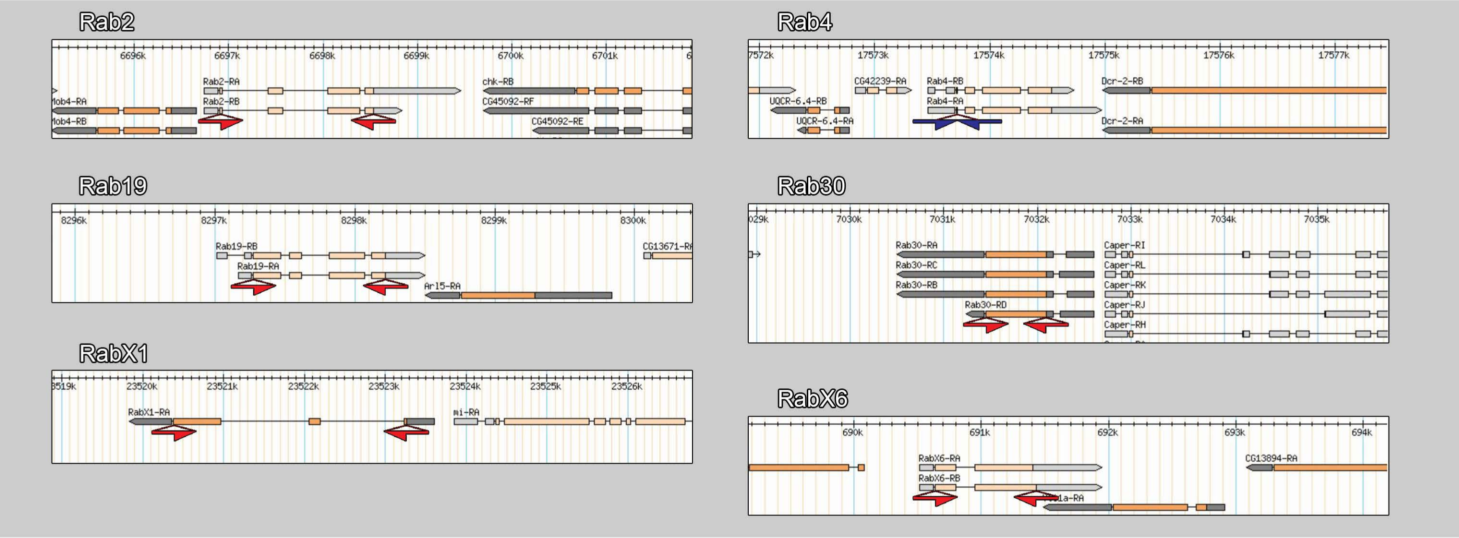


C CRISPR knock-in cassette

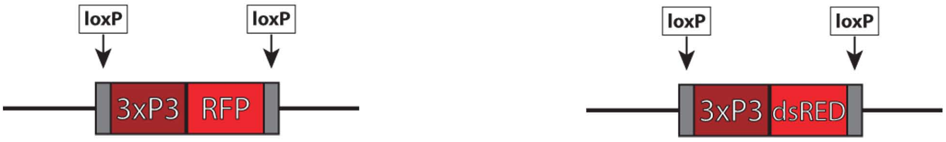


D CRISPR mutants

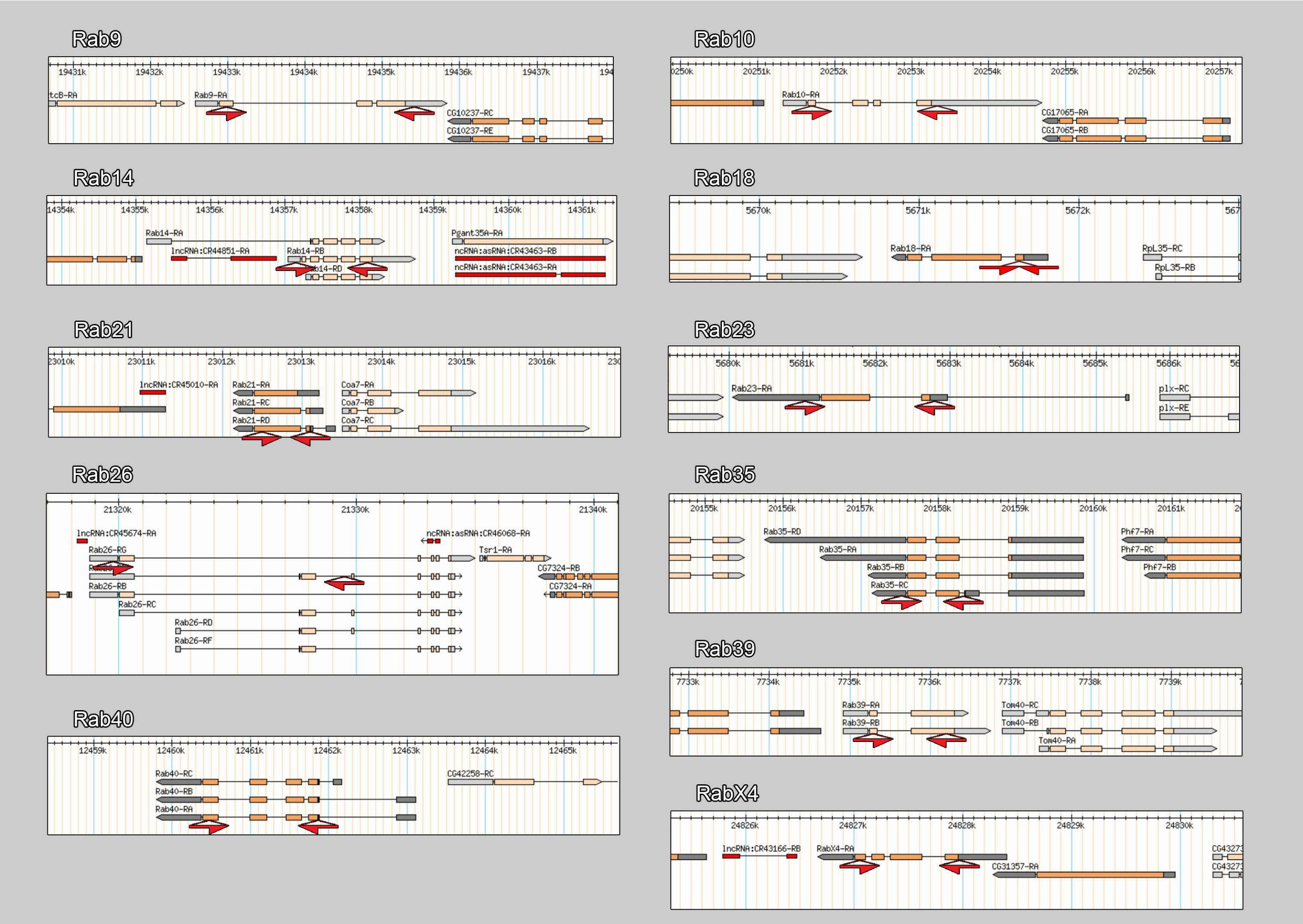

### Supplementary Figure 2

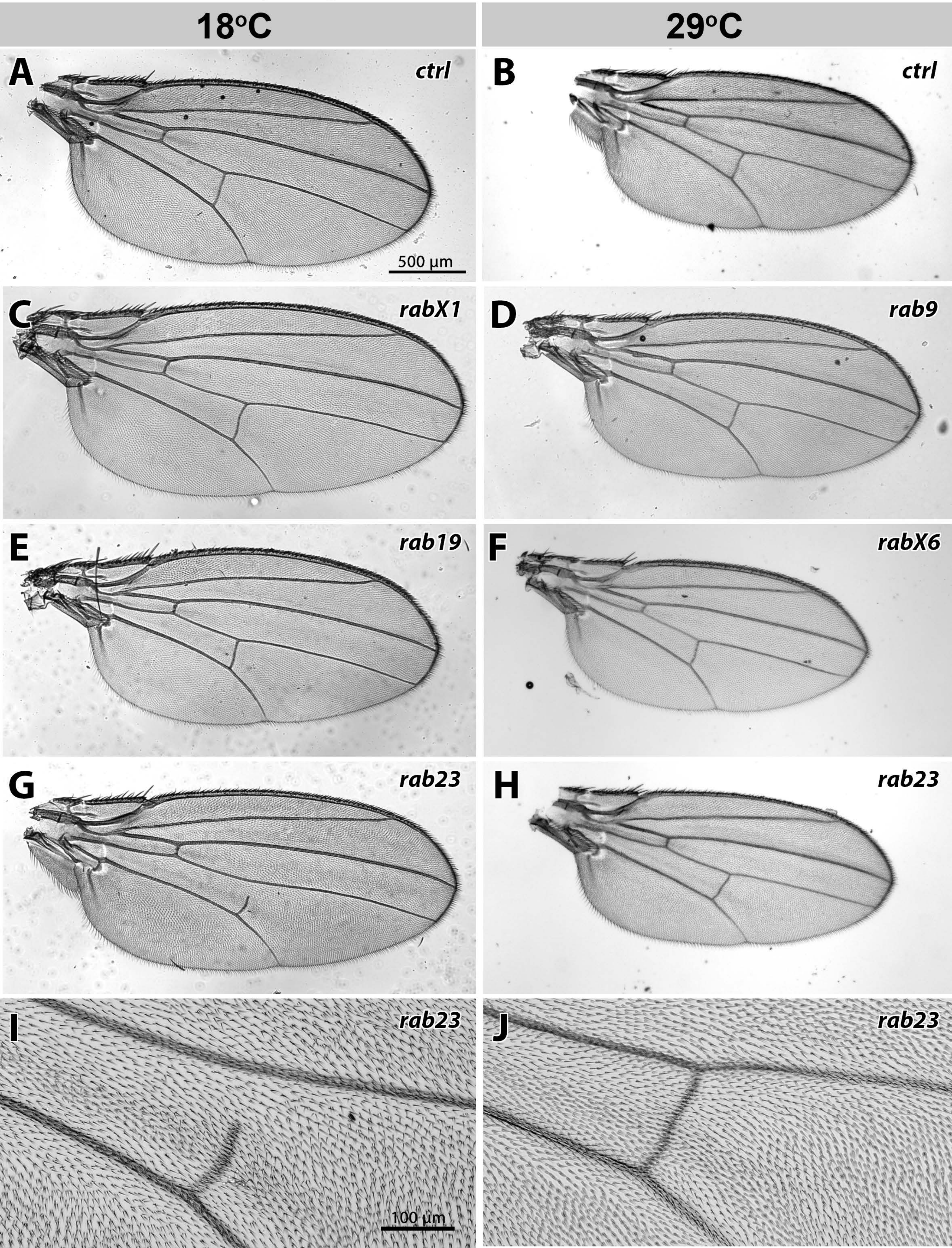

### Supplementary Figure 3

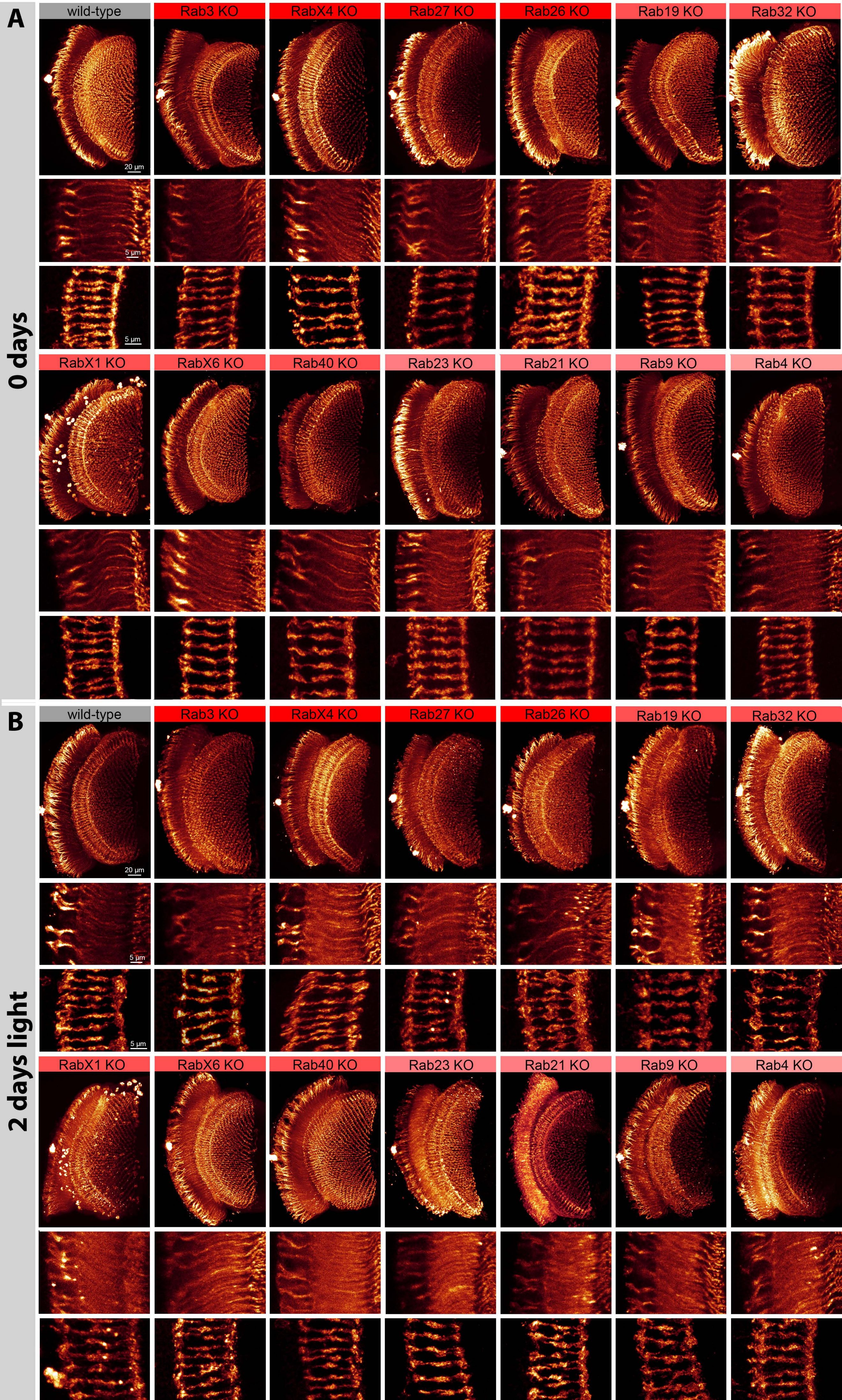

### Supplementary Figure 4

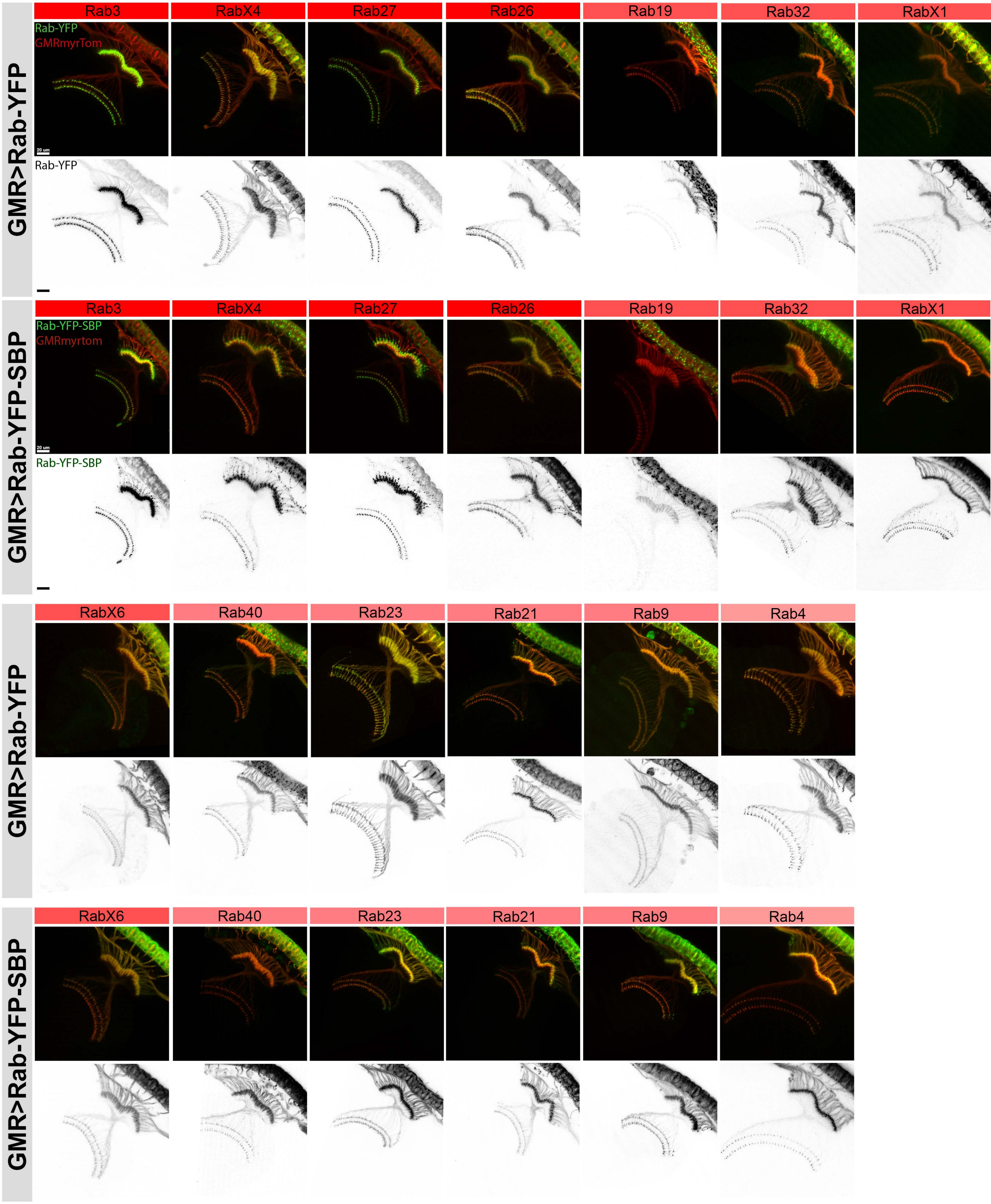

### Supplementary Figure 5

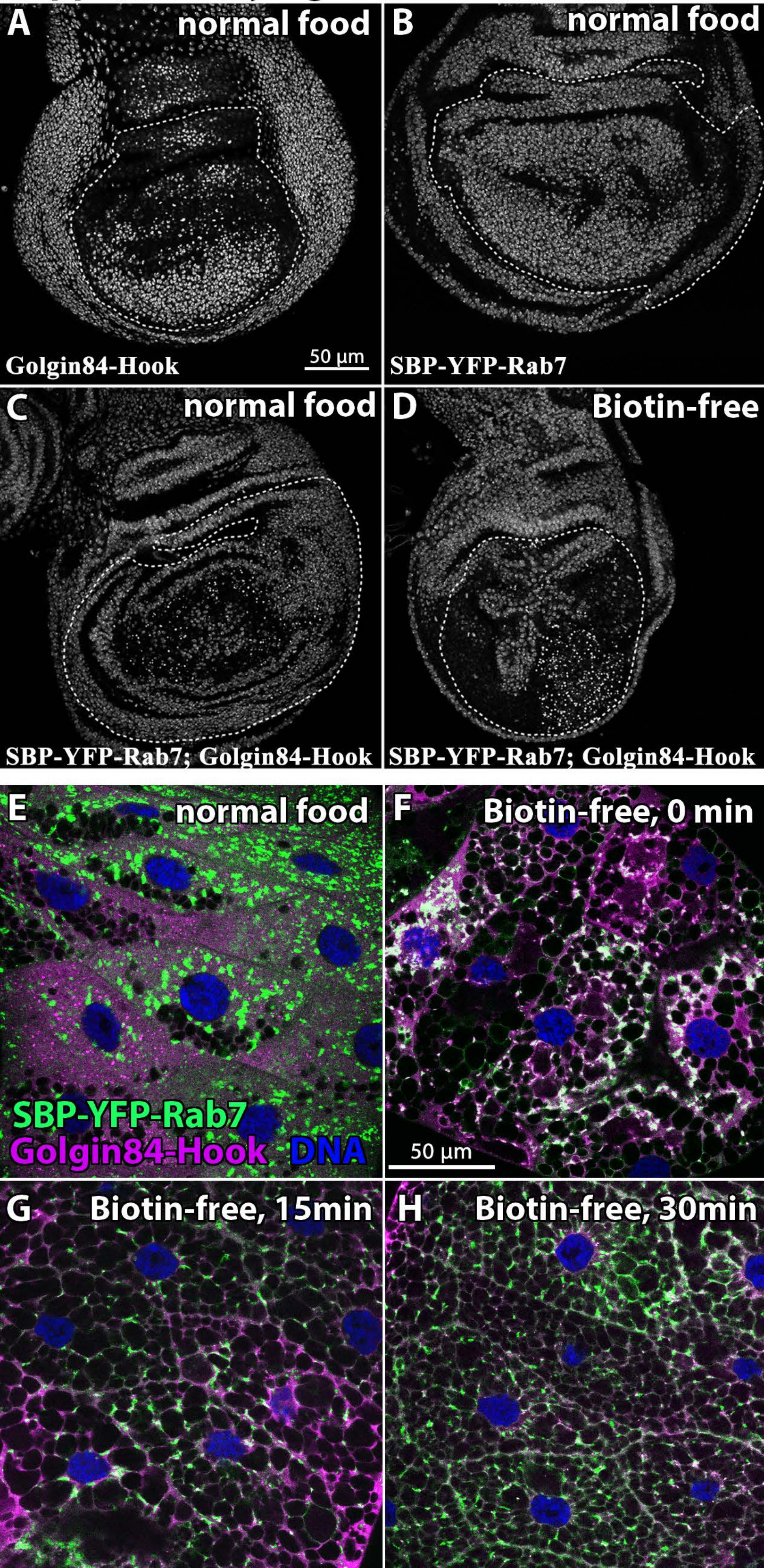

**Supplementary Figure 6**

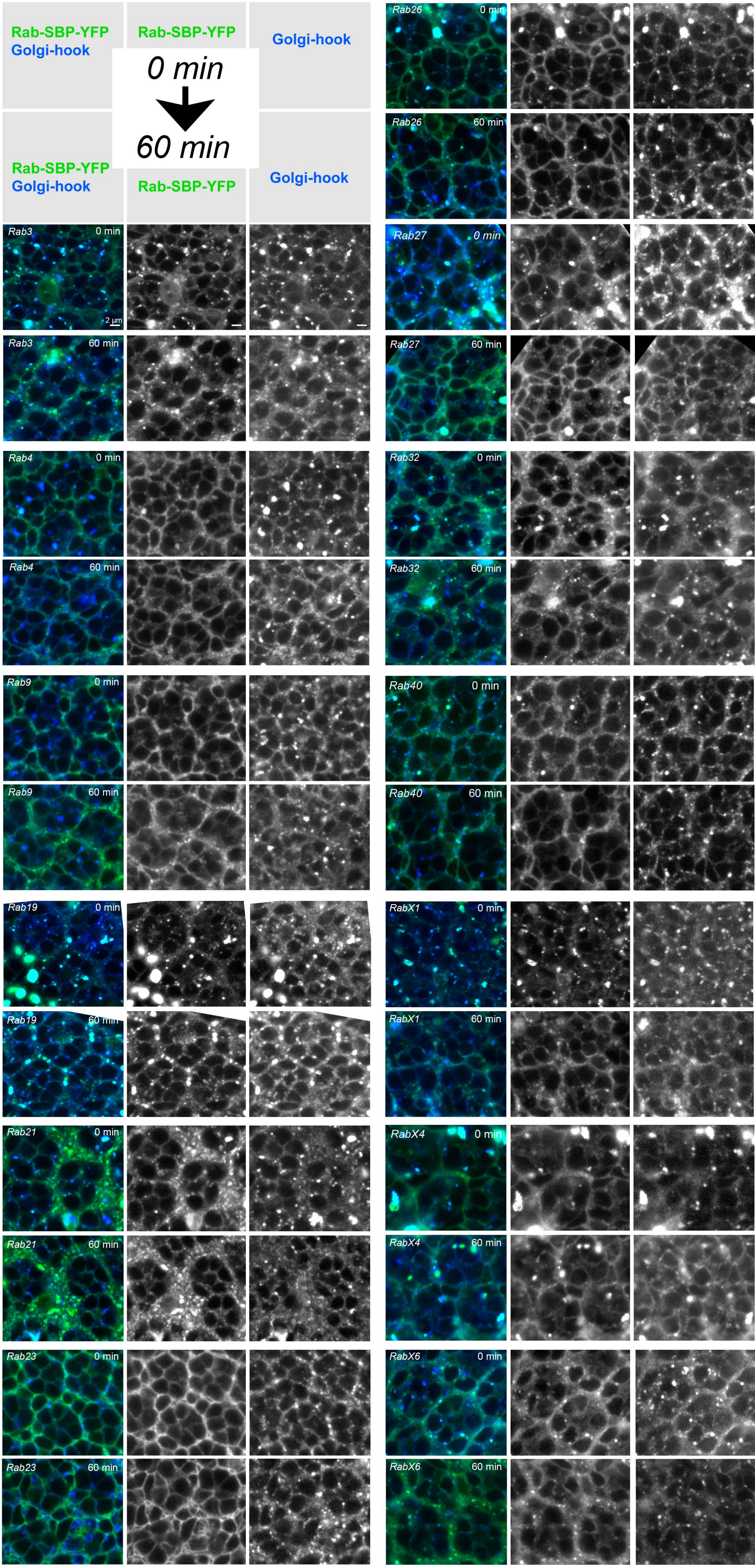

### Supplementary Figure 7

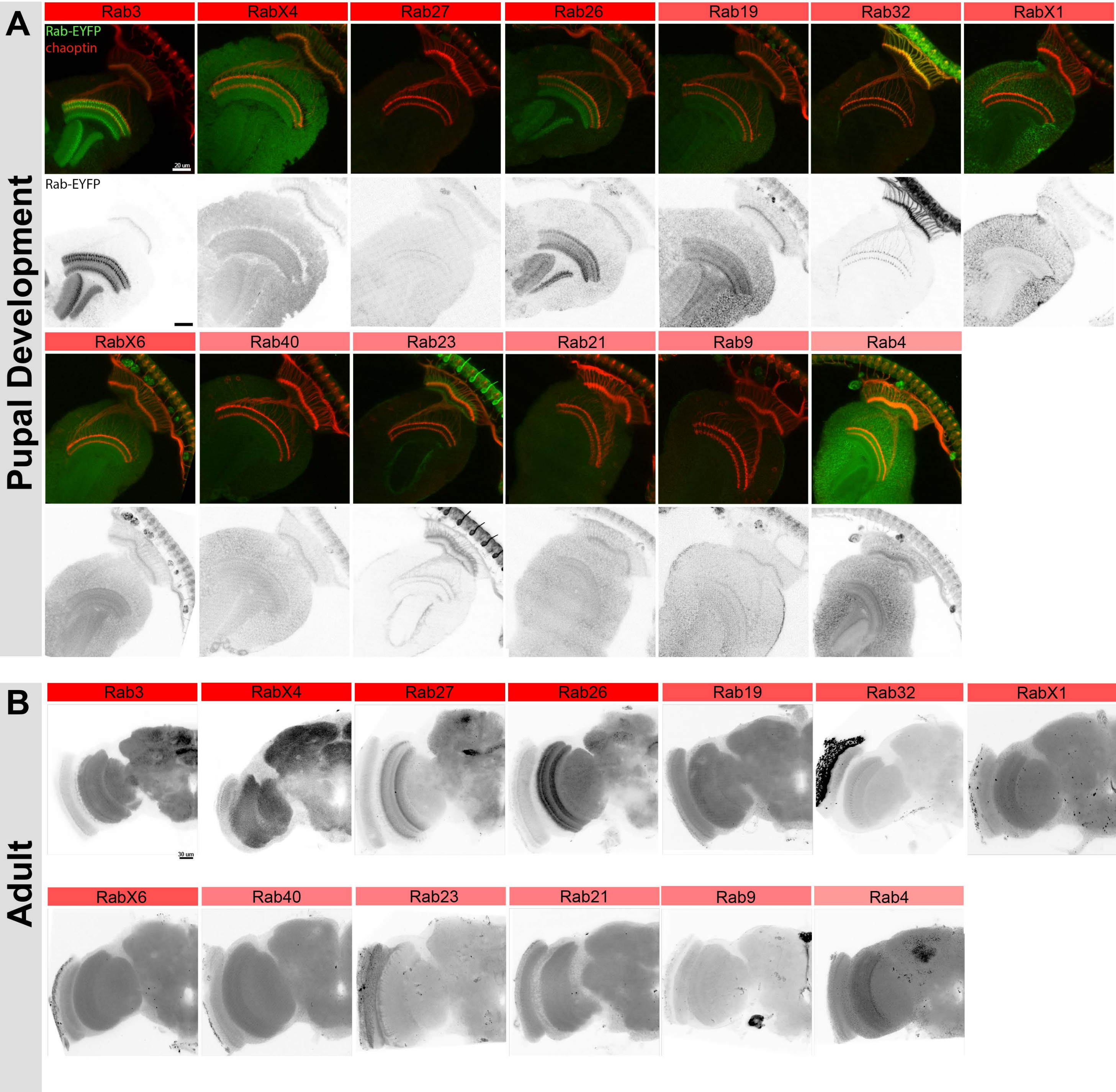

#### Supplementary Table 1

| 18 degree | days until… (after egg collection) | | | number of adults in total | Ø number of adults per vial |
|---|---|---|---|---|---|
| | first instar larva | pupa | adult | | |
| control | 0 | 8.43 ± 0.13 | 16.8 ± 0.14 | 170 | 8.1 ± 1.82 |
| *rab3* | 0.4 ± 0.13 | 9.29 ± 0.18 | 18.1 ± 0.24 | 159 | 7.57 ± 1.02 |
| *rab4* | 0.1 ± 0.07 | 9.19 ± 0.09 | 17.38 ± 0.11 | 516 | 24.57 ± 2.1 |
| *rab9* | 0.29 ± 1 | 7.86 ± 0.08 | 16.71 ± 0.12 | 344 | 16.38 ± 1.41 |
| *rab14* | 0.1 ± 0.07 | 7.57 ± 0.11 | 15.76 ± 0.1 | 473 | 22.52 ± 1.82 |
| *rab18* | 0.33 ± 0.13 | 9.05 ± 0.21 | 17.62 ± 0.25 | 203 | 9.67 ± 1.17 |
| *rab19* | 0.1 ± 0.07 | 8.76 ± 0.14 | 18.1 ± 0.17 | 275 | 13.1 ± 1.44 |
| *rab21* | 0.29 ± 0.12 | 8.29 ± 0.1 | 16.95 ± 0.16 | 368 | 17.52 ± 0.98 |
| *rab23* | 0.24 ± 0.1 | 8.62 ± 0.11 | 17.19 ± 0.11 | 239 | 11.38 ± 1.01 |
| *rab26* | 0.24 ± 0.1 | 8.71 ± 0.1 | 17 ± 0.1 | 442 | 21.05 ± 1.91 |
| *rab27* | 0.1 ± 0.07 | 8.86 ± 0.1 | 17.1 ± 0.07 | 368 | 17.52 ± 1.16 |
| *rab32* | 0.1 ± 0.07 | 9.38 ± 0.11 | 18 ± 0.17 | 316 | 15.05 ± 0.87 |
| *rab39* | 0 | 9.05 ± 0.13 | 17.76 ± 0.1 | 523 | 24.9 ± 1.75 |
| *rab40* | 0.33 ± 0.16 | 10.19 ± 0.09 | 18.67 ± 0.14 | 261 | 12.42 ± 0.98 |
| *rabX1* | 1.71 ± 0.46 | 10.56 ± 0.26 | 19.24 ± 0.2 | 37 | 1.76 ± 0.32 |
| *rabX4* | 0.9 ± 0.14 | 12.5 ± 0.3 | 21 ± 0.53 | 8 | 0.38 ± 0.13 |
| *rabX6* | 0 | 9.14 ± 0.14 | 17.8 ± 0.13 | 446 | 21.24 ± 2.64 |

| 25 degree | days until… (after egg collection) | | | number of adults in total | Ø number of adults per vial |
|---|---|---|---|---|---|
| | first instar larva | pupa | adult | | |
| control | 0.15 ± 0.08 | 4.86 ± 0.08 | 8.95 ± 0.05 | 230 | 10.95 ± 2.01 |
| *rab3* | 0.2 ± 0.09 | 5 ± 0.12 | 9.2 ± 0.12 | 197 | 9.85 ± 1.45 |
| *rab4* | 0.1 ± 0.07 | 5.29 ± 0.1 | 9.25 ± 0.16 | 634 | 30.19 ± 2.06 |
| *rab9* | 0.14 ± 0.08 | 5 | 9 | 411 | 19.57 ± 1.63 |
| *rab14* | 0.1 ± 0.07 | 4.62 ± 0.11 | 8.67 ± 0.14 | 499 | 23.76 ± 1.75 |
| *rab18* | 0.05 ± 0.05 | 4.86 ± 0.08 | 8.9 ± 0.07 | 345 | 16.43 ± 1.34 |
| *rab19* | 0.24 ± 0.1 | 4.95 ± 0.08 | 9.71 ± 0.1 | 379 | 18.04 ± 1.51 |
| *rab21* | 0.19 ± 0.09 | 5 | 8.95 ± 0.05 | 412 | 19.62 ± 1.74 |
| *rab23* | 0.1 ± 0.07 | 4.86 ± 0.08 | 9 | 305 | 14.52 ± 1.71 |
| *rab26* | 0.2 ± 0.09 | 4.95 ± 0.05 | 9.1 ± 0.07 | 480 | 22.86 ± 2.1 |
| *rab27* | 0.14 ± 0.08 | 5 | 9 | 382 | 18.19 ± 1.12 |
| *rab32* | 0.29 ± 0.1 | 5.24 ±0.1 | 9.19 ± 0.11 | 459 | 21.86 ± 1.62 |
| *rab39* | 0.24 ± 0.1 | 5.14 ± 0.08 | 9.43 ± 0.11 | 627 | 29.86 ± 2.69 |
| *rab40* | 0.24 ± 0.1 | 5.7 ± 0.13 | 9.95 ± 0.15 | 241 | 11.48 ± 1.14 |
| *rabX1* | 0.56 ± 0.15 | 5.29 ± 0.14 | 9.67 ± 0.84 | 40 | 1.9 ± 0.35 |
| *rabX4* | 0.89 ± 0.08 | 7.14 ± 0.7 | 12 ± 0.85 | 1 | 0.05 ± 0.05 |
| *rabX6* | 0.05 ± 0.05 | 5.05 ± 0.05 | 9.23 ± 0.1 | 459 | 21.86 ± 2.62 |

| 29 degree | days until… (after egg collection) | | | # number of adults in total | Ø number of adults per vial |
|---|---|---|---|---|---|
| | first instar larva | pupa | adult | | |
| control | 0.15 ± 0.09 | 4 | 7.67 ± 0.11 | 112 | 5.33 ± 1.21 |
| *rab3* | 0.1 ± 0.07 | 4.19 ± 0.09 | 7.57 ± 0.11 | 168 | 8 ± 0.81 |
| *rab4* | 0 | 4.52 ± 0.11 | 8 | 572 | 27 ± 2.14 |
| *rab9* | 0.14 ± 0.08 | 3.95 ± 0.05 | 7.57 ± 0.11 | 369 | 17.57 ± 1.73 |
| *rab14* | 0 | 4.05 ± 0.05 | 7.43 ± 0.11 | 516 | 24.57 ± 1.66 |
| *rab18* | 0.1 ± 0.07 | 4.19 ± 0.09 | 7.35 ± 0.11 | 291 | 13.86 ± 1.68 |
| *rab19* | 0.19 ± 0.09 | 4.05 ± 0.05 | 8.94 ± 0.18 | 132 | 7.33 ± 1.4 |
| *rab21* | 0.05 ± 0.05 | 4.19 ± 0.09 | 8 ± 0.2 | 378 | 18.9 ± 1.72 |
| *rab23* | 0.29 ± 0.1 | 4 | 7.95 ± 0.05 | 288 | 13.71 ± 0.92 |
| *rab26* | 0.29 ± 0.1 | 4.14 ± 0.08 | 7.81 ± 0.09 | 378 | 18 ± 1.76 |
| *rab27* | 0.05 ± 0.05 | 4 | 7.43 ± 0.11 | 444 | 21.14 ± 0.95 |
| *rab32* | 0.24 ± 0.1 | 4.33 ± 0.11 | 8 | 307 | 14.62 ± 1.01 |
| *rab39* | 0.19 ± 0.09 | 4.1 ± 0.07 | 7.81 ± 0.09 | 572 | 27.24 ± 2.3 |
| *rab40* | 0.33 ± 0.11 | 4 | 7.33 ± 0.33 | 280 | 13.33 ± 0.74 |
| *rabX1* | 0.81 ± 0.16 | 4.75 ± 0.11 | 8.19 ± 0.1 | 38 | 1.81 ± 0.38 |
| *rabX4* | 0.75 ± 0.1 | 6.38 ± 0.56 | 13.5 ± 3.5 | 2 | 0.1 ± 0.07 |
| *rabX6* | 0.19 ± 0.09 | 4.05 ± 0.05 | 7.76 ± 0.1 | 445 | 21.19 ± 2.75 |
